# UNC-30/PITX coordinates neurotransmitter identity with postsynaptic GABA receptor clustering

**DOI:** 10.1101/2024.02.14.580278

**Authors:** Edgar Correa, Morgane Mialon, Mélissa Cizeron, Jean-Louis Bessereau, Berangere Pinan-Lucarre, Paschalis Kratsios

**Affiliations:** Department of Neurobiology, University of Chicago, Chicago, IL, 60637, USA; Committee on Cell and Molecular Biology, University of Chicago, Chicago, IL, 60637, USA; University of Chicago Neuroscience Institute, Chicago, IL, 60637, USA; Melis, Universite Claude Bernard Lyon 1, CNRS UMR5284, INSERM U1314, Institut NeuroMyoGene - Faculte de Medecine et de Pharmacie, 69008 Lyon, France

**Author notes:** equal contribution (Edgar Correa, Morgane Mialon).

## Abstract

Terminal selectors are transcription factors that control neuronal identity by regulating the expression of key effector molecules, such as neurotransmitter (NT) biosynthesis proteins, ion channels and neuropeptides. Whether and how terminal selectors control neuronal connectivity is poorly understood. Here, we report that UNC-30 (PITX2/3), the terminal selector of GABA motor neuron identity in *C. elegans*, is required for NT receptor clustering, a hallmark of postsynaptic differentiation. Animals lacking *unc-30* or *madd-4B,* the short isoform of the MN-secreted synapse organizer *madd-4* (*Punctin/ADAMTSL*), display severe GABA receptor type A (GABA_A_R) clustering defects in postsynaptic muscle cells. Mechanistically, UNC-30 acts directly to induce and maintain transcription of *madd-4B* and GABA biosynthesis genes (e.g., *unc-25/GAD*, *unc-47/VGAT*). Hence, UNC-30 controls GABA_A_R clustering on postsynaptic muscle cells and GABA biosynthesis in presynaptic cells, transcriptionally coordinating two critical processes for GABA neurotransmission. Further, we uncover multiple target genes and a dual role for UNC-30 both as an activator and repressor of gene transcription. Our findings on UNC-30 function may contribute to our molecular understanding of human conditions, such as Axenfeld-Rieger syndrome, caused by PITX2 and PITX3 gene mutations.

## INTRODUCTION

In the nervous system, neuronal communication critically depends on the proper transmission of signals through chemical and electrical synapses. In the context of chemical synapses, presynaptic neurons must be able to synthesize and package into synaptic vesicles specific chemical substances known as neurotransmitters (NTs), such as acetylcholine (ACh), gamma-aminobutyric acid (GABA), and glutamate (Glu). Upon secretion into the synaptic cleft, each NT molecule binds to its cognate receptors located at the postsynaptic cell membrane, thereby evoking postsynaptic electrical responses.

Genes encoding proteins for NT biosynthesis and packaging (e.g., enzymes, transporters) are co-expressed in specific neuron types. The co-expression of these proteins defines the NT identity (or NT phenotype) of individual neuron types (e.g., cholinergic, GABAergic, dopaminergic). Although instances of NT identity switching have been described in the nervous system^1,2^, it is generally the case that individual neuron types acquire a specific NT identity during development and maintain it throughout life. The continuous expression of NT identity genes is fundamental for the ability of a presynaptic neuron to signal to its postsynaptic targets. For efficient neurotransmission, however, it is equally important that cognate NT receptors cluster at postsynaptic domains precisely juxtaposed to presynaptic boutons ^3,4^. Whether and how these two critical processes, i.e., NT identity of the presynaptic neuron and NT receptor clustering at the postsynaptic cell, are coordinated remains poorly understood.

Genetic studies in *C. elegans*, flies and mice have revealed a phylogenetically conserved principle for the control of NT identity: neuron type-specific transcription factors, termed “terminal selectors”, coordinate the expression of NT identity genes, thereby coordinating synthesis of enzymes and transporters necessary for NT biosynthesis and signaling^5,6^. In addition, terminal selectors broadly control batteries of genes encoding proteins essential for neuronal identity and function (e.g., ion channels, neuropeptides)^6,7^. To date, terminal selectors have been described for 111 of the 118 *C. elegans* neuron types ^8,9^. Beyond *C. elegans*, terminal selectors have also been identified in fruit flies (*Drosophila)*, cnidarians (*Nematostella vectensis)*, marine chordates (*Ciona intestinalis)*, and mice (*Mus musculus*)^6^, suggesting a deeply conserved role for these critical regulators of NT identity. A defining feature of terminal selectors is their continuous expression - from development throughout adulthood - in specific neuron types^5^. While the essential roles of terminal selectors in establishing NT identity during development is well-attested across model organisms, their involvement in maintaining NT identity in later life stages remains poorly examined^10^, partially due to the lack of genetic tools for inducible terminal selector depletion in late life stages.

In the case of GABAergic neurons, NT identity is defined by the co-expression of highly conserved proteins, including (a) the enzyme glutamic acid decarboxylase (GAD) which synthesizes GABA from its precursor, (b) the vesicular GABA transporter (VGAT) which packages GABA into synaptic vesicles, and (c) the GABA re-uptake transporter (GAT) (**Figure 1**)^12^. Importantly, reduced expression of these GABA identity determinants, as well as impaired GABA transmission, lead to a variety of neuropsychiatric diseases, including schizophrenia, autism, epilepsy or anxiety ^13^ ^14^.

**Figure 1:**
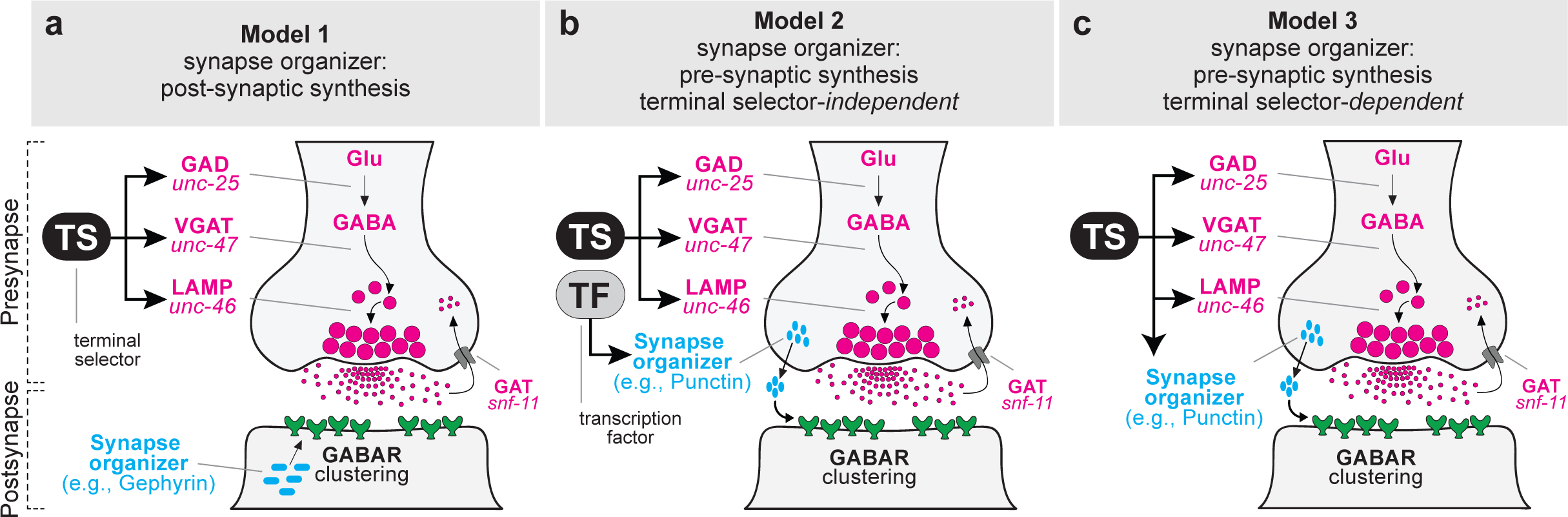
Three hypothetical models for transcriptional control of GABA synapse organizers. (a-c) A detailed description of the three models is provided in the Introduction.

Despite GABA being the most abundant inhibitory NT both in invertebrate and vertebrate nervous systems, it is poorly understood how the expression of GABA identity genes is controlled over time, from development through adulthood, to ensure GABA neurotransmission. To date, a handful of studies in *C. elegans* and mice have identified terminal selectors in various GABAergic neuron types. Examples include homeodomain proteins (e.g., UNC-30/PITX)^15–18^, nuclear hormone receptors (e.g., NHR-67/NR2E1)^12^, and GATA-type (GATA2/3) transcription factors, each necessary for expression of GABA identity genes during development ^19–21^. However, whether any of these factors is required for maintaining GABA identity gene expression during post-embryonic life is unknown^10^.

During neuronal development, GABA receptor (GABAR) clustering is fundamental for postsynaptic differentiation, a process primarily driven by synapse-organizing molecules that can be either secreted or bound to the cell membrane ^13^. In mice, the cell adhesion molecule Neuroligin-2, the scaffold protein Gephyrin, and the transmembrane protein β-dystroglycan act as synapse organizers to control GABAR clustering^22–24^. In the nematode *C. elegans*, the secreted molecule MADD-4 (Muscle Arm Development Defect-4), also known as human Punctin/*ADAMTSL,* acts as an anterograde synapse organizer at neuromuscular synapses ^25^. Specifically, the short MADD-4 isoform (MADD-4B) activates DCC (Deleted in Colorectal Cancer)/UNC-40 signaling, recruiting an intracellular postsynaptic scaffold composed of FRM-3, a FERM (p4.1, Ezrin, Radixin, Moesin) domain protein, and LIN-2/CASK (CAlcium Calmodulin dependent Serine/threonine Kinase)^26,27^. Moreover, MADD-4B controls GABAR positioning at synapses by recruiting the sole *C. elegans* neuroligin homolog, NLG-1, which binds to LIN-2^26–29^. On the other hand, the long *madd-4* isoform, MADD-4L, promotes ACh receptor (AChR) clustering on muscle cells through the formation of an extracellular scaffold^25,30–32^.

In vertebrates, there are two *madd-4* orthologs: *Punctin1/Adamtsl1* and *Punctin2/Adamtsl3* ^33,34^. Although the brain function of *Punctin1/Adamtsl1* remains unknown, recent data identified Adamtsl3 as an extracellular synapse organizer in the rodent hippocampus, where it supports glutamatergic and GABAergic synapse formation *in vivo*^35^. Furthermore, in the adult brain, Adamtsl3 signals via DCC at GABAergic synapses and facilitates synapse maintenance, synaptic plasticity, and memory. In humans, *Adamtsl3* is widely expressed in the brain, and has been identified as a candidate gene for schizophrenia^36^. Despite their well-established roles in GABAR clustering, the transcriptional mechanisms that control expression of the aforementioned synapse organizers remain poorly understood.

GABA neurotransmission critically relies on (a) the ability of the presynaptic neuron to continuously express of GABA identity genes (e.g., GAD, VGAT, GAT) and (b) the ability of the postsynaptic neurons to properly cluster GABARs ^3,4^. Whether these two processes, that occur in two synaptically-connected cells, are coordinated remains poorly understood. In principle, at least three non-mutually exclusive models can be envisioned for the transcriptional control of a GABA synapse organizer (**Figure 1**). GABAR clustering at the post-synaptic (target) cell could be achieved via the activity of a synaptic organizer (membrane-bound or secreted) produced in the post-synaptic cell. For example, Gephyrin, a synapse organizer produced in the target cell, is essential for GABAR clustering (**Figure 1a**, model 1). Alternatively, GABA-R clustering in the post-synaptic cell may rely on secreted synaptic organizers, like MADD-4/Punctin, produced in the presynaptic GABAergic neuron (**Figure 1b-c**, models 2 - 3). In that case, transcription of the synapse organizer gene may or may not require the activity of the terminal selector of the presynaptic neuron (model 2 versus 3). Our previous work in *C. elegans* provided support for model 3 in the context of cholinergic neuromuscular synapses^37^; the terminal selector UNC-3 (Collier, Ebf) is not only required for AChR clustering in the postsynaptic neuron, but also controls NT identity genes in the presynaptic cell. However, whether this principle of transcriptional coordination extends beyond cholinergic motor neurons (MNs) was unclear.

*C. elegans* has been a prime model to dissect molecular mechanisms underlying NT identity and synapse formation^4,8^. Here, we show that the *C. elegans* terminal selector of GABAergic MN identity, UNC-30/PITX, is required for clustering of type A GABARs, a major type of inhibitory NT receptors^38,39^. We find that UNC-30 acts directly to activate transcription of the synapse organizer *madd-4B*. Hence, the terminal selector UNC-30 coordinates GABAR clustering on postsynaptic muscle cells (via control of *madd-4B/Punctin*) with acquisition of GABAergic identity in presynaptic MNs (**Figure 1**, model 3). Further, we find that UNC-30 acts directly to maintain the expression of *madd-4B* and NT identity genes (e.g., *unc-25*/GAD, *unc-47*/VGAT) in late larval and adult stages. Intriguingly, UNC-30 also represses transcription of the long *madd-4* isoform (*madd-4L)*, which is normally required for AChR clustering in postsynaptic muscle cells^25^. Hence, our work in GABA MNs highlights that NT receptor clustering, a central event of postsynaptic differentiation, is transcriptionally coordinated with acquisition and maintenance of NT identity, significantly extending previous observations made in *C. elegans* cholinergic MNs to other neuron types^37^. Last, we uncovered additional target genes that are either positively or negatively regulated by UNC-30/PITX, indicating direct activator and repressor functions. Such mechanistic insights may help us understand the molecular mechanisms underlying human genetic disorders caused by *PITX* gene mutations, such as Axenfeld-Rieger syndrome^40–42^.

## RESULTS

### The experimental system: GABAergic neuromuscular synapses in *C. elegans*

*C. elegans* locomotion relies on both cholinergic and GABAergic MNs, whose cell bodies intermingle along the ventral nerve cord (equivalent to vertebrate spinal cord) (**Figure 2a**). Based on anatomical criteria, cholinergic and GABAergic MNs are respectively divided into six (VA, VB, DA, DB, AS, VC) and two (DD, VD) classes, which form *en passant* synapses along the ventral and dorsal nerve cords (**Figure 2a**)^4,43^. The coordinated activity of excitatory cholinergic and inhibitory GABAergic MNs generates sinusoidal locomotion in *C. elegans,* with each muscle cell receiving dual innervation from cholinergic and GABAergic MNs. Along the dorsal nerve cord of adult animals, three cholinergic MN classes (DA, DB, and AS) form dyadic synapses, providing excitatory input not only to dorsal muscles but also to VD GABAergic neurons, which in turn innervate and inhibit ventral muscles (**Figure 2a**)^44^. Along the ventral nerve cord, another three cholinergic MN classes (VA, VB, and VC) also form dyadic synapses with ventral muscles and DD GABAergic neurons, which innervate and inhibit dorsal muscles (**Figure 2a**). Because each muscle cell receives both excitatory (ACh) and inhibitory (GABA) inputs, the *C. elegans* neuromuscular system represents a powerful model to study how different NT receptors precisely cluster in front of their corresponding neurotransmitter release sites^4^.

**Figure 2:**
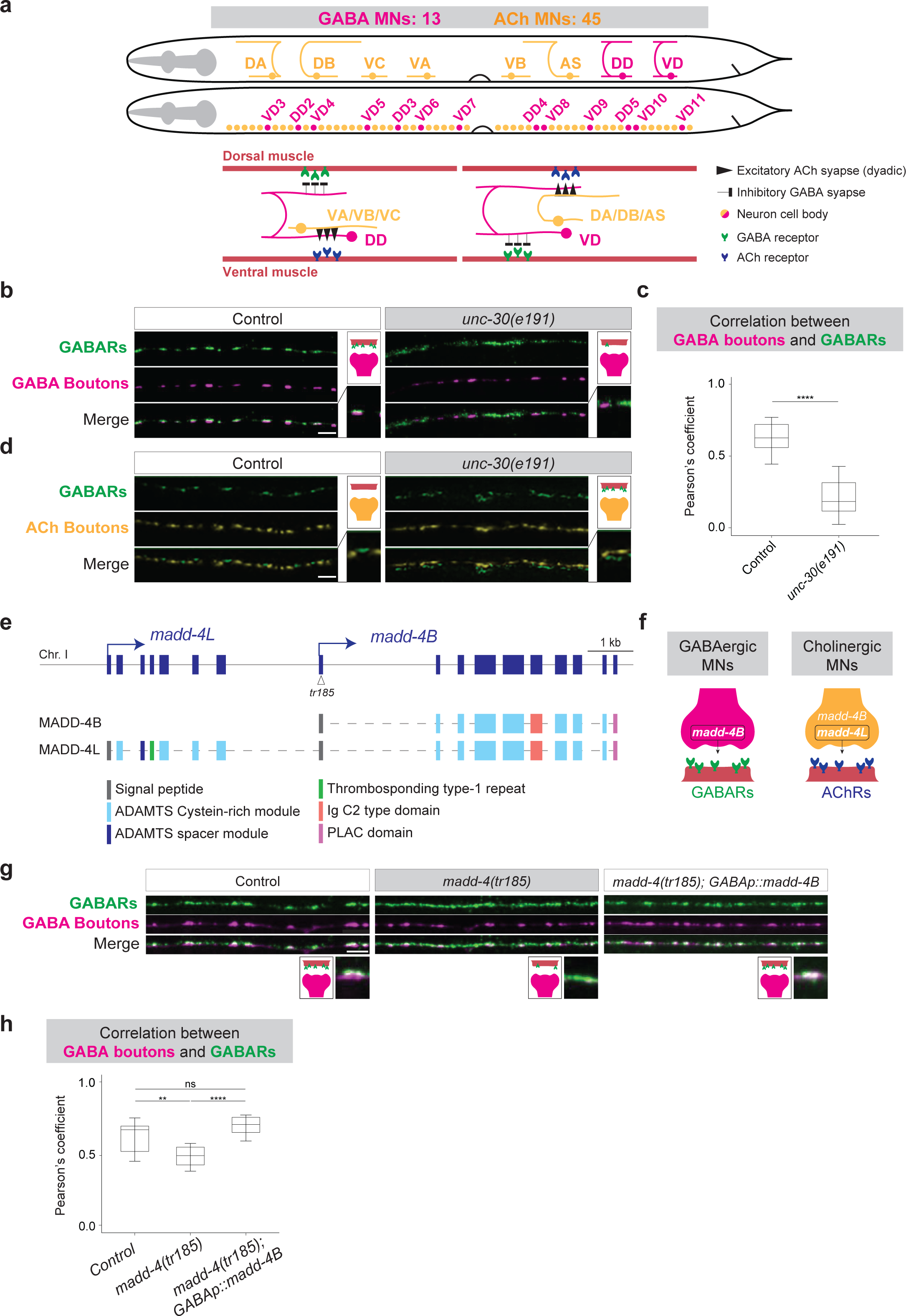
*unc-30* and *madd-4B* control GABAR clustering at *C. elegans* neuromuscular synapses. (a) MN connectivity in the *C. elegans* ventral nerve cord. Cholinergic (DA, DB, VC, VA, VB, AS) and GABAergic (DD, VD) MN cell bodies intermingle. See text for details. (b) Fluorescence micrographs of GABA_A_R (*kr296 [unc-49::rfp],* green pseudocolor) and GABAergic presynaptic boutons (*otEx5663 [unc-30prom::GFP::rab-3],* magenta pseudocolor) in control and *unc-30(e191)* animals. (c) Pearson’s correlation coefficient between GABA_A_R and GABAergic presynaptic boutons as shown in (b). Box and whisker plots show median, lower, and upper quartiles – whiskers represent SD. ANOVA followed by Tukey post-hoc test. ****p<0.0001. Control: n=11, *unc-30(e191)*: n=10. (d) Immunofluorescence staining of GABA_A_R (anti-UNC-49 antibody) and cholinergic presynaptic boutons (anti-UNC-17 antibody) in control and *unc-30(e191)* animals. (e) *madd-4* locus with both *(madd-*4*L*, *madd-4B*) isoforms. Protein domains are shown. The *madd-4(tr185)* allele carries a mutation in exon 1 of *madd-4B*. (f) Schematic of *madd-4B* and *madd-4L* expression in *C. elegans* MNs. (g) Fluorescence micrographs of GABA_A_R (*kr296[unc-49::rfp],* green pseudocolor) and GABAergic boutons (*krIs67[unc-47prom::SNB-1::BFP]*, magenta pseudocolor) in control and *madd-4(tr185)* animals, and a rescue strain: *madd-4B* in GABAergic MNs (*krSi92 [unc-47prom::T7::MADD-4S::GFP]*. Scale bar: 5 µm. (h) Pearson’s correlation coefficient between GABA_A_R and GABAergic presynaptic boutons as shown in (g). Kruskal-Wallis followed by Dunn’s post-test. ns: non-significant, *p < 0.01, ****p < 0.0001. Wild-type: n=18; *madd-4(tr185):* n=22; rescue strain: n=21.

### *unc-30/PITX* controls GABA_A_ receptor clustering at inhibitory neuromuscular synapses in a non-cell autonomous manner

Within the *C. elegans* ventral nerve cord, the transcription factor UNC-30/PITX2-3 is specifically expressed in GABAergic (DD, VD) MNs^17^. In these cells, UNC-30 controls the expression of GABA identity genes (**Figure 1**) ^15–18,45^. Recent studies also implicated UNC-30 in synaptic remodeling, as it prevents premature synapse rewiring of DD cells and aberrant synapse rewiring of VD cells^46,47^. However, whether UNC-30 is necessary for the postsynaptic differentiation of target muscle cells remained unknown.

We therefore asked whether genetic loss of *unc-30* affects GABAR clustering in *C. elegans* muscle cells innervated by GABAergic MNs. To this end, we used an endogenous *RFP* reporter for *unc-*49 (UNC-49::RFP), which encodes a type-A GABAR (GABA_A_R) expressed both in ventral and dorsal body wall muscles (**Figure 2b**)^27^. Upon crossing this strain to a presynaptic marker (*unc-30prom::GFP::RAB-3)* for GABAergic MNs (DD, VD), we visualized in young adult (day 1) animals the juxtaposition of GABAergic presynaptic boutons and GABA_A_R clusters in body wall muscles (**Figure 2b**). We focused our analysis on DD synapses onto dorsal muscle (**Figure 2a**) because the dorsal nerve cord (DNC) does not contain MN cell bodies, thereby facilitating the visualization of UNC-49::RFP and *unc-30prom::GFP::RAB-3* fluorescent signals (**Figure 2a-b**).

In homozygous animals carrying a strong loss-of-function *unc-30* allele (*e191*)^17,48^, we found that GABA_A_Rs are present on dorsal muscle (DNC), but no longer cluster opposite presynaptic GABA (DD) boutons at young adult (day 1) stages (**Figure 2b-c**). Because in control animals GABAergic (DD) and cholinergic (DA, DB, AS) neurons form *en passant* neuromuscular synapses with dorsal muscle (**Figure 2a**), we considered two possibilities: either GABA-Rs on dorsal muscle of *unc-30(e191)* animals are not juxtaposed to any presynaptic terminal, or they are inappropriately juxtaposed to presynaptic boutons of cholinergic (DA, DB, AS) MNs. We therefore performed double immunofluorescence staining against UNC-49 and UNC-17 (VAChT/SLC18A3), a marker of cholinergic presynaptic boutons. We indeed found that GABA_A_Rs incorrectly localize opposite to cholinergic presynaptic boutons in the *unc-30(e191)* mutants (**Figure 2d**). Therefore, *unc-30/PITX* is necessary for the correct positioning of GABA_A_Rs at neuromuscular synapses along the dorsal nerve cord. Because *unc-30* is present in GABAergic MNs but not expressed in body wall muscles or muscle progenitor cells (**Supplementary Figure 1**) ^17^, we conclude that *unc-30* controls GABA_A_R clustering in an indirect (non-cell autonomous) manner.

### The short isoform of *madd-4 (Punctin)* controls GABA_A_R clustering at neuromuscular synapses in a non-cell autonomous manner

In a previous study, we demonstrated that *madd-4/Punctin*, a secreted synapse organizer, is critical for GABAR and AChR clustering at *C. elegans* neuromuscular synapses^25^. The *madd-4* locus generates two isoforms through alternative promoter usage (**Figure 2e**)^37,49^. The long isoform (*madd-4L*) is produced by cholinergic MNs and required for levamisole-sensitive AChR (L-AChR) clustering at neuromuscular synapses (**Figure 2e-f**)^25,37^. The short isoform (*madd-4B*) is required for GABAR clustering (**Figure 2e-f**)^25^. Because *madd-4B* is produced by both GABAergic and cholinergic MNs^25,26,37^, it remained unclear whether *madd-4B* from GABAergic and/or cholinergic MNs is required for GABA_A_ receptor clustering at neuromuscular synapses.

To test this, we first analyzed animals specifically lacking *madd-4B* gene activity using the *madd-4(tr185)* allele (**Figure 2e**). Confirming their previously reported synaptic phenotype^26,27^, we found that UNC-49::RFP fluorescence signal on the dorsal muscle of *madd-4B(tr185)* animals is no longer restricted to sites opposite of GABA (DD) boutons (**Figure 2g-h**). Instead, UNC-49::RFP is detected along the dorsal nerve cord. Because both GABAergic and cholinergic neuromuscular synapses are located *en passant,* the continuous distribution of UNC-49::RFP along the dorsal nerve cord suggests that GABA_A_R clusters face both GABAergic (DD) and cholinergic (DA, DB, AS) presynaptic boutons (**Figure 2a**). Importantly, expression of *madd-4B* specifically in GABAergic MNs led to a complete rescue of this phenotype (**Figure 2g-h**). Because *madd-4B* is expressed in GABAergic neurons, but not in muscle cells^25,37,49^, these findings consolidate a non-cell autonomous role for the secreted synaptic organizer *madd-4B* in GABA_A_R clustering at postsynaptic muscle cells.

### UNC-30 controls *madd-4B* transcription in GABAergic MNs

Because both *unc-30* and *madd-4B* mutants display defects in GABA_A_R localization (**Figure 2c, 2h**), we hypothesized that the transcription factor UNC-30 regulates *madd-4B* in GABAergic MNs. To test this, we employed CRISPR/Cas9 genome editing and generated an endogenous fluorescent reporter of *madd-4B* transcription. Before the ATG of *madd-4B*, we inserted a *mScarlet* sequence preceded by two copies of a nuclear localization signal (2xNLS) and followed by the SL2 trans-splicing element (**Figure 3a**). The *2x::NLS::mScarlet* sequence and endogenous *madd-4B* are transcribed as one mRNA, but each is translated independently due to the SL2 element. In agreement with transgenic *madd-4B* reporters^37^, this endogenous *2xNLS::mScarlet::SL2::madd-4B* transcriptional reporter (*mScarlet::madd-4B* hereafter) is expressed both in cholinergic and GABAergic MNs, albeit higher levels are observed in GABAergic MNs (**Figure 3b**). To test the effect of *unc-30* gene loss in *madd-4B* expression specifically in GABAergic MNs, we crossed a nuclear marker for cholinergic MNs (*cho-1::SL2::YFP::H2B*) to the *mScarlet::madd-4B* reporter in the context of control and *unc-30(e191)* animals (**Figure 3b**). We observed a significant decrease in the number of GABAergic cells (defined by the absence of *cho-1::SL2::YFP::H2B* signal) expressing *mScarlet::madd-4B* in *unc-30(e191)* mutants at the fourth larval (L4) stage (**Figure 3b-c**). That is, all 13 GABAergic neurons (DD2-DD5, VD3-VD11) of the ventral nerve cord express *mScarlet::madd-4B* in control animals, but only ∼10 neurons in *unc-30(-)* mutants (**Figure 3a, c**). Importantly, the remaining *mScarlet::madd-4B* expression in these 10 GABAergic neurons is also decreased, as revealed by quantification of *mScarlet::madd-4B* fluorescence intensity with single-cell resolution (e.g., VD3, DD2, VD4, VD5, DD3, VD6) (**Figure 3d**). The remaining *madd-4B* expression suggests that additional, yet-to-be-identified factors cooperate with UNC-30 to activate *madd-4B* expression in these cells. We note that throughout our analysis we excluded six GABAergic MNs (DD1, DD6, VD1, VD2, VD13) because their location (outside the ventral nerve cord) makes their identification less straightforward.

**Figure 3:**
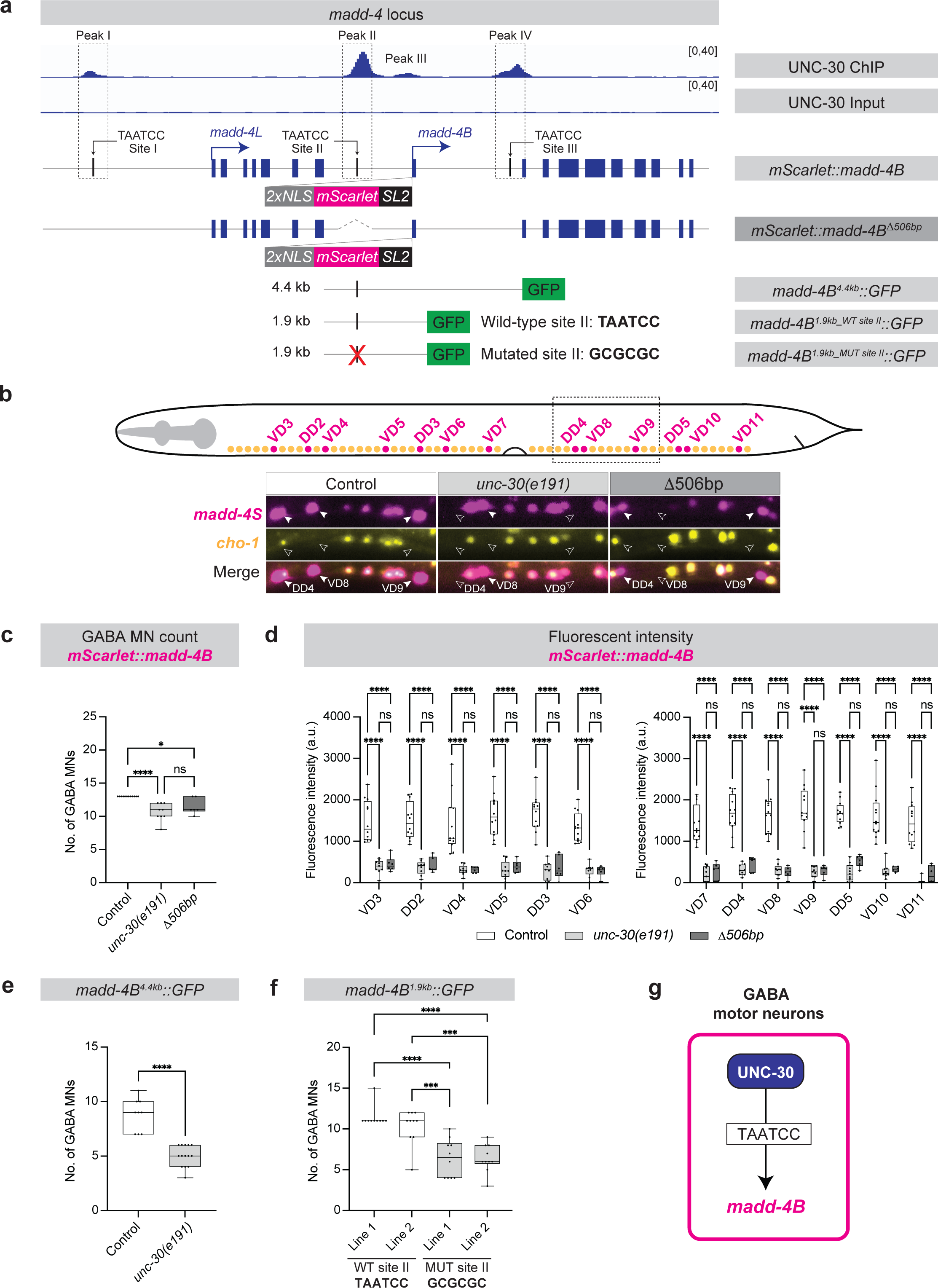
UNC-30 directly activates *madd-4B* in GABAergic MNs. (a) UNC-30 ChIP-Seq and Input (negative control) on *madd-4* locus. Four UNC-30 binding peaks (peaks I, II, III, IV) and three UNC-30 binding sites (TAATCC sites I, II, II) are shown. Depicted below: endogenous *madd-4B* reporter *(syb623[2xNLS::mScarlet::SL2::madd-4B])*, mutant reporter allele (*syb3561[2xNLS::mScarlet::SL2:: madd-4B^Δ^*^506b^*^p^]*), *madd-4B* transgenic reporters: *(otEx5601 [madd-4B*^4^.^4k^*^b^::GFP], otEx4948-9[madd-4B*^1^.^9k^*^b^::GFP]), kasEx315-6[madd-4B*^1^.^9k^*^b_GCGCGC::^GFP*]). (b) GABAergic (magenta) and cholinergic (yellow) MNs. Dashed box depicts imaged area. Fluorescence micrographs of *madd-4B (syb623[2xNLS::mScarlet::SL2::madd-4B])* and a cholinergic MN reporter *(otIs354[cho-1(fosmid)::SL2::YFP::H2B])* in control and *unc-30(e191)* animals, and *syb3561[2xNLS::mScarlet::SL2:: madd-4B^Δ^*^506b^*^p^]*) animals. GABA MNs: *mScarlet+;YFP –;* cholinergic MNs: *mScarlet +;YFP+*. Images of day 1 adults. White arrowheads: GABAergic MNs. (c) Quantification of number of GABAergic MNs of animal genotypes shown in (b). Unpaired t-test with Welch’s correction. ****p<0.0001. Control: n=10, *unc-30(e191)*: n=10, Δ506bp mutant: n=8. (d) Quantification of *madd-4B (2xNLS::mScarlet::SL2::madd-4B)* fluorescent intensity in individual GABAergic MNs. Two-way ANOVA followed by Sidak’s multiple comparison test. ^ns^p>0.05, *p<0.002, ****p<0.0001. Wild-type: n=10, *unc-30(e191:* n=10, Δ506bp mutant: n=8. (e) Quantification of GABAergic MNs expressing *madd-4B(otEx5601[madd-4B*^4^.^4k^*^b^::GFP])* in control and *unc-30(e191)* animals. Unpaired t-test with Welch’s correction. ^ns^p>0.05, ****p<0.0001. Control: n=9, *unc-30(e191)*: n=14. (f) Quantification of GABAergic MNs expressing *otEx4948-9[madd-4B*^1^.^9k^*^b_TAATCC^::GFP]* and *kasEx315-6[madd-4B*^1^.^9k^*^b_GCGCGC^::GFP]s.* One-way ANOVA followed by Sidak’s multiple comparison test. ***p<0.001, ****p<0.0001. Wild-type line1: n=10, wild-type line 2: n=10, *madd-4B*^1^.^9k^*^b_GCGCGC^* line 1: n=10, *madd-4B*^1^.^9k^*^b_GCGCGC^* line 2: n=10. (g) Schematic of UNC-30 directly activating *madd-4B*. For panels c-f: Box and whisker plots show median, lower, and upper quartiles – whiskers represent minimum and maximum. Black circles depict values.

The single-cell resolution of our analysis indicates that *unc-30* controls *madd-4B* both in DD (e.g., DD2, DD3) and VD (e.g., VD3, VD4, VD5) neurons (**Figure 3d**). Corroborating this observation, we quantified *mScarlet::madd-4B* expression at larval stage 1 (L1), a developmental stage at which only DD (not VD) neurons are present in the *C. elegans* nerve cord (**Supplementary Figure 2a**). Again, we found a significant decrease in *madd-4B* expression in DD neurons of *unc-30(e191)* mutants (**Supplementary Figure 2b**). In agreement with our endogenous transcriptional reporter (*mScarlet::madd-4B*), expression of a transgenic translational *madd-4B* reporter is also affected in *unc-30* animals at L1^50^. Altogether, we conclude that *unc-30* controls endogenous *madd-4B* transcription in GABAergic MNs, and this effect is observed both at early (L1) and late (L4) larval stages.

### UNC-30 directly activates *madd-4B* transcription in GABAergic MNs

Because *madd-4B* expression is reduced in GABA MNs of *unc-30(e191)* animals (**Figure 3b-d**), we investigated whether *madd-4B* is a direct target of UNC-30. Leveraging an available dataset of chromatin immunoprecipitation followed by sequencing (ChIP-Seq)^51^, we identified UNC-30 binding at four genomic regions (peaks I-IV): Peak I is upstream of *madd-4L*, whereas peaks II-IV surround the first exon of *madd-4B* (**Figure 3a**). Within peaks I, II, and IV, we identified a canonical UNC-30 binding site (TAATCC)^15,16^. To test whether UNC-30 binding upstream of *madd-4B* is required for *madd-4B* expression, we employed CRISPR/Cas9 genome editing to delete a 506 bp-long region that spans peak II (Δ506bp, **Figure 3a**). This manipulation was conducted in animals carrying the endogenous *mScarlet::madd-4B* reporter. Similar to *unc-30(e191)* mutants, we observed a decrease in the number of GABAergic MNs expressing *mScarlet* in L4 stage animals homozygous for the 506bp deletion (**Figure 3b-c**), as well as a decrease in the levels of *mScarlet* expression in individual GABAergic MNs (**Figure 3d**).

ChIP-Seq data and our analysis of *mScarlet::madd-4B^Δ^*^506b^*^p^* animals strongly indicate that UNC-30 acts directly to activate *madd-4B* transcription. To further test direct transcriptional control, we examined transgenic animals carrying different transcriptional reporters of *madd-4B* (**Figure 3a**). First, we found that reporters containing DNA sequences either 4.4kb (*madd-4B*^4^.^4k^*^b^::GFP*) or 1.9kb (*madd-4B*^1^.^9k^*^b^::GFP*) upstream of *madd-4B* (both containing peak II) drive *GFP* expression in GABA MNs (**Figure 3a, e, f**), consistent with the endogenous *madd-4B::mScarlet* reporter (**Figure 3c**). Second, *madd-4B*^4^.^4k^*^b^::GFP* reporter expression depends on *unc-30*, evidenced by a reduction in the number of GABA MNs expressing *GFP* in *unc-30(e191)* mutants (**Figure 3e**). Third, we found that mutation of the UNC-30 binding site II (wild type: TAATCC, mutated: GCGCGC) results in a significant decrease in the number of GABA MNs expressing *madd-4B*^1^.^9k^*^b^::GFP* (**Figure 3f**). Altogether, we conclude that UNC-30 acts directly to activate *madd-4B* transcription in GABA MNs (**Figure 3g**).

### UNC-30 represses *madd-4L* transcription in GABAergic MNs

The ChIP-Seq data also showed UNC-30 binding (peak I) in the *cis*-regulatory region upstream of exon 1 of *madd-4L* (**Figure 3a**). Because *madd-4L* is known to be specifically expressed in cholinergic MNs^25,37^, we hypothesized that UNC-30 binds directly upstream of *madd-4L* to repress its transcription in GABA MNs. Again, we employed CRISPR/Cas9 genome editing and generated an endogenous *mScarlet* reporter for *madd-4L* (*2x::NLS::mScarlet::SL2::madd-4L*), referred hereafter as *mScarlet::madd-4L* (**Figure 4a**). Supporting our hypothesis, we observed ectopic expression of *mScarlet::madd-4L* in GABA MNs of *unc-30(e191)* mutant animals both at L1 (**Supplementary Figure 3**) and at L4 (**Figure 4b-d**). We found that up to 13 GABA MNs of the ventral nerve cord ectopically express *mScarlet::madd-4L* in *unc-30(e191)* mutants (**Figure 4b-d**).

**Figure 4:**
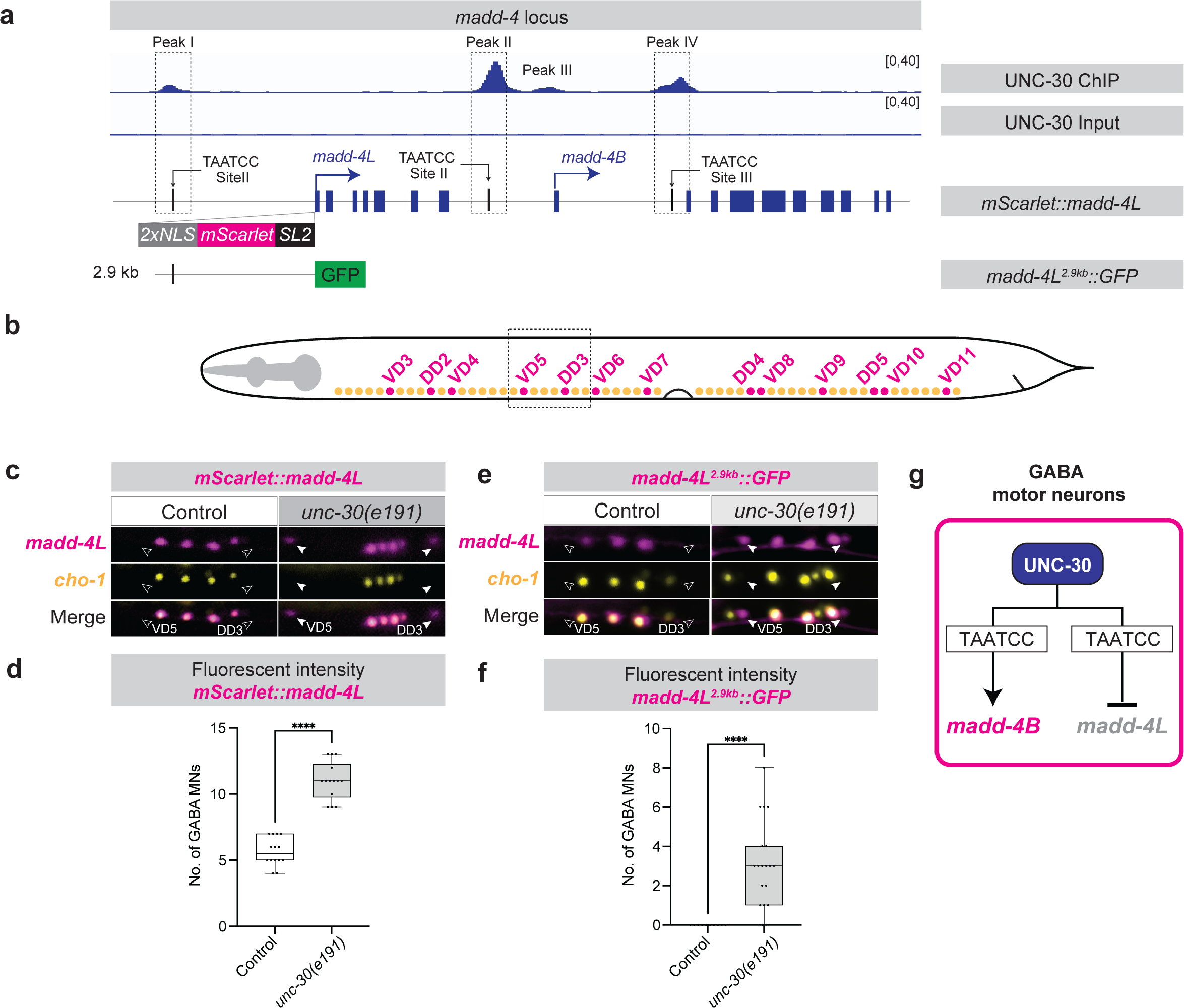
UNC-30 represses *madd-4L* in GABAergic motor neurons. (a) UNC-30 ChIP-Seq and Input (negative control) tracks on *madd-4* locus. Depicted below: (1) endogenous (*syb624[2xNLS::mScarlet::SL2::madd-4L]*) and 2) transgenic (*sEx14990 [madd-4L(2.9kb prom)::GFP]*) *madd-4L* reporter. (b) GABAergic (magenta) and cholinergic (yellow) MNs in *C. elegans*. Dashed box depicts imaged area shown in (c and e). (c) Fluorescence micrographs of *madd-4L*(*syb624 [2xNLS::mScarlet::SL2::madd-4L]*) and a cholinergic motor neuron reporter *(otIs354[cho-1(fosmid)::SL2::YFP::H2B])* in control and *unc-30(e191)* animals. GABAergic MNs: *mScarlet+;YFP–*, cholinergic MNs: *mScarlet+;YFP+*. Images of day1 adults. White arrowheads: GABAergic MNs. (d) Quantification of GABAergic MNs expressing *madd-4L(syb624 [2xNLS::mScarlet::SL2::madd-4L])* as shown in (c). Unpaired t-test with Welch’s correction. ****p<0.0001. Wild-type: n=13, *unc-30(e191)*: n=13. (e) Fluorescence micrographs of *madd-L*(*sEx14990[madd-4L(2.9kb prom)::GFP])* and a cholinergic MN reporter*(otIs544 [cho-1(fosmid)::SL2::mCherry::H2B])* in control and *unc-30(e191)* animals. GABAergic MNs: *GFP+; mCherry -*–; cholinergic MNs: *GFP+;mCherry+*. Images of day 1 adults. White arrowheads: GABAergic MNs. (f) Quantification of the number of GABAergic MNs expressing *madd-4L*(*sEx14990 [madd-4L(2.9kb prom)::GFP])* as shown in (e). Unpaired t-test with Welch’s correction. ****p<0.0001. Wild-type: n=13, *unc-30(e191)*: n=13. (g) Schematic: dual role of UNC-30 in controlling *madd-4* isoforms. For panels d and f: Box and whisker plots show median, lower, and upper quartiles – whiskers represent minimum and maximum. Black circles depict values.

UNC-30 binding (peak I) upstream of *madd-4L* suggests UNC-30 directly represses *madd-4L* in GABAergic MNs. In agreement with this notion, a transcriptional *madd-4L*^2^.^9k^*^b^::GFP* reporter driving *GFP* under the control of a 2.9kb *cis*-regulatory region upstream of *madd-4L* (contains peak I) shows ectopic expression in GABA MNs of *unc-30* mutant animals at L4 (**Figure 4a, 4e-f**). We conclude that, in GABA MNs, UNC-30/PITX controls two isoforms of the same synapse organizer in opposite ways; it directly activates *madd-4B* and represses *madd-4L* (**Figure 4g**).

### UNC-30 is continuously required to maintain *madd-4B* expression in GABAergic MNs

The continuous expression of both *unc-30* and *madd-4B* in GABAergic MNs, from larval stages throughout adulthood, raises the question of whether UNC-30 is required continuously to activate *madd-4B* expression. We therefore generated an inducible *unc-30* allele, leveraging the auxin-inducible degradation (AID) system ^53,54^. Using CRISPR/Cas9, we introduced the *mNG::3xFLAG::AID* cassette before the *unc-30* STOP codon (**Figure 5a**). The resulting *unc-30::mNG::3xFLAG::AID* allele *(syb2344)* serves as an endogenous fluorescent (mNG, mNeonGreen) reporter of the UNC-30 protein, which can be degraded upon auxin treatment due to the presence of the AID degron (**Figure 5b**). We generated double homozygous animals for *unc-30::mNG::3xFLAG::AID* and *ieSi57 (Peft-3::TIR1::mRuby),* the latter providing pansomatic expression of *TIR1* – an F-box protein that binds to AID in the presence of auxin, leading to proteasomal degradation of UNC-30::mNG::3xFLAG::AID. As proof-of-principle, we first assessed UNC-30::mNG::3xFLAG::AID levels in individual GABA MNs in ethanol-treated (control) or 4mM auxin-treated animal for 2 days, from L3 and to adult day 1 (**Figure 5b**). Compared to ethanol-treated animals, auxin-treated animals showed a robust reduction in the levels of UNC-30::mNG::3xFLAG::AID fluorescent intensity, indicating efficient depletion (**Figure 5c-e**). Auxin-treated animals exhibited a mild reduction in the total number of *mScarlet::madd-4B-expressing* MNs (**Figure 5f-g**), but a significant reduction in *mScarlet* fluorescent intensity levels in all individual GABA MNs (**Figure 5f-h, Supplementary Figure 4**). We therefore conclude that UNC-30 is required during late larval and young adult stages to maintain *madd-4B* expression in GABAergic MNs (**Figure 5k**). UNC-30’s continuous requirement is likely critical to maintain GABA_A_R clustering throughout life.

**Figure 5:**
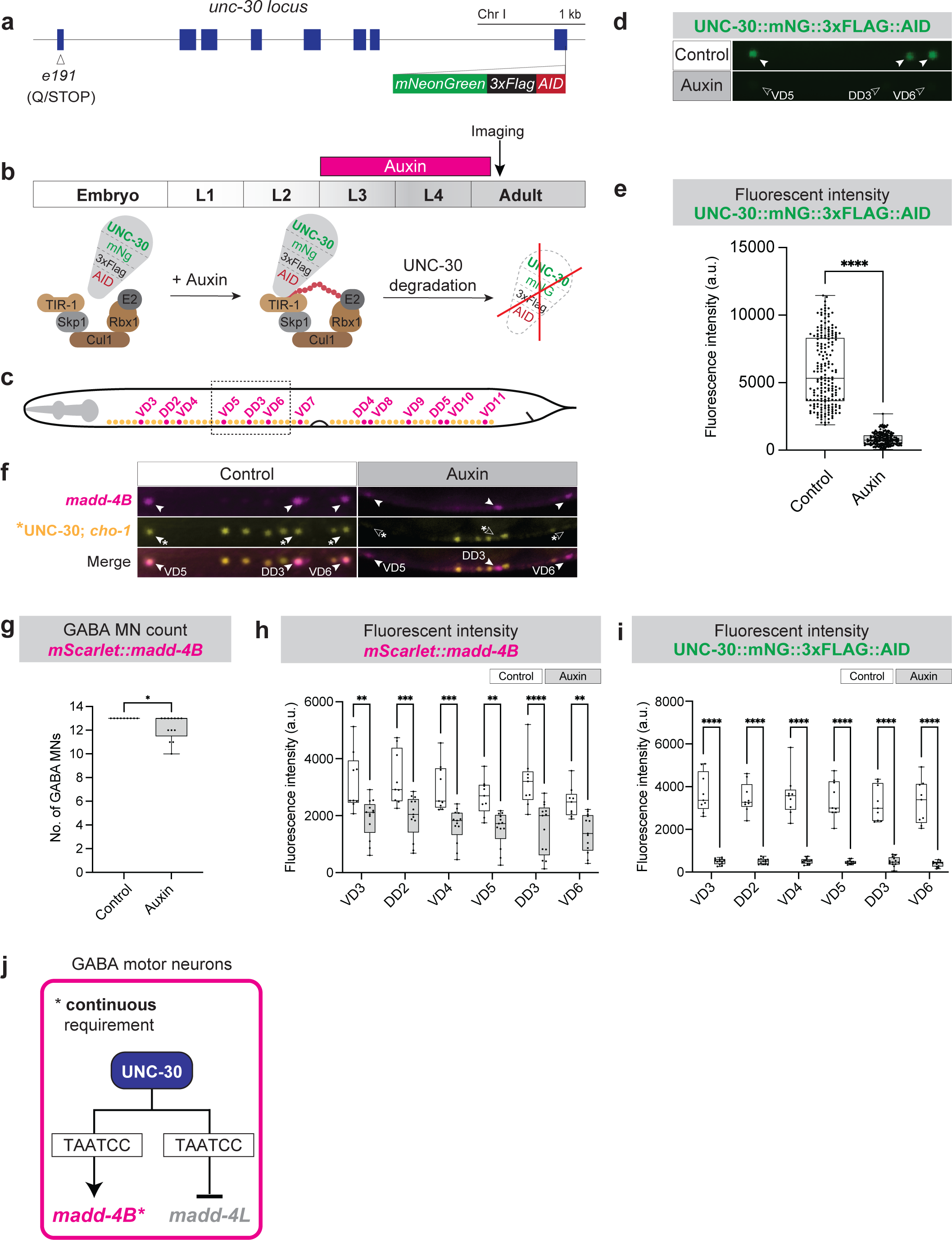
UNC-30 is required to maintain *madd-4B* expression in GABA MNs. (a) Schematic of *unc-30* locus and *unc-30::mNG::3xFLAG::AID* allele (*syb2344*). (b) Schematic of AID system. The E3 ligase complex: Skp1, Cul1, Rbx1, E2. Auxin treatment from L3 to adult day 1 stage. Imaging occurred at day 1. (c) GABAergic (magenta) and cholinergic (yellow) MNs. Dashed box depicts imaged area shown in (d,f). (d-e) Fluorescence micrographs of UNC-30 (*syb2344 [UNC-30::mNG::3xFLAG::AID])* in control (EtOH treated) and auxin treated animals. Animals express TIR1 pan-somatically (*ieSi57 [Peft-3::TIR1::mRuby]*). White arrowheads: GABAergic MNs. Quantification of UNC-30 (*syb2344 [UNC-30::mNG::3xFLAG::AID]*fluorescent intensity in GABAergic MNs. Unpaired t-test with Welch’s correction. ****p<0.0001. Control: n=195, Auxin-treated=195. (f) Fluorescence micrographs of *madd-4B (syb623[2xNLS::mScarlet::SL2::madd-4B]),* UNC-30 (*syb2344[UNC-30::mNG::3xFLAG::AID])*, and a cholinergic MN reporter *(otIs354[cho-1(fosmid)::SL2::YFP::H2B])* in control (EtOH treated) and auxin-treated animals. White arrowheads: GABAergic MNs. Green asterisk: GABA MNs expressing UNC-30::mNG::3xFLAG::AID. (g) Quantification of the number of GABAergic MNs expressing *madd-4B (2xNLS::mScarlet::SL2::madd-4B)* as shown in (f). Unpaired t-test with Welch’s correction. *p<0.01. Wild-type: n=9, *unc-30(e191)*: n=13. (h-i) Quantification of (h) *madd-4B (2xNLS::mScarlet::SL2::madd-4B)* or (i) UNC-30 (*syb2344 [UNC-30::mNG::3xFLAG::AID])* fluorescent intensity in GABAergic MNs, as shown in (f). Two-way ANOVA followed by Sidak’s multiple comparison test. **p<0.002, ***p<0.0002, ****p<0.0001. Control: n=9, Auxin-treated=13. (j) UNC-30 is required to maintain *madd-4*.. In panels e, h, and i: Box and whisker plots show median, lower, and upper quartiles – whiskers represent minimum and maximum. Black circles depict values.

### UNC-30 is required to maintain expression of GABA biosynthesis genes

Prompted by our *madd-4B* observations, we next asked whether UNC-30 is continuously required to maintain the expression of additional target genes. A previous study employing a constitutive null allele (*e191*) showed that UNC-30 activates the expression of two GABA identity genes during development, *unc-25*/GAD and *unc-47*/VGAT ^16^. Mutating the UNC-30 binding site (TAATCC) in transgenic *unc-25* and *unc-47* reporter animals resulted in reduced reporter expression in GABA MNs, strongly suggesting UNC-30 regulates these targets via direct binding^16^. Consistent with these previous findings, analysis of the UNC-30 ChIP-Seq dataset showed UNC-30 binding in the *cis*-regulatory regions of *unc-25* and *unc-47* endogenous loci (**Figure 6c, f**).

**Figure 6:**
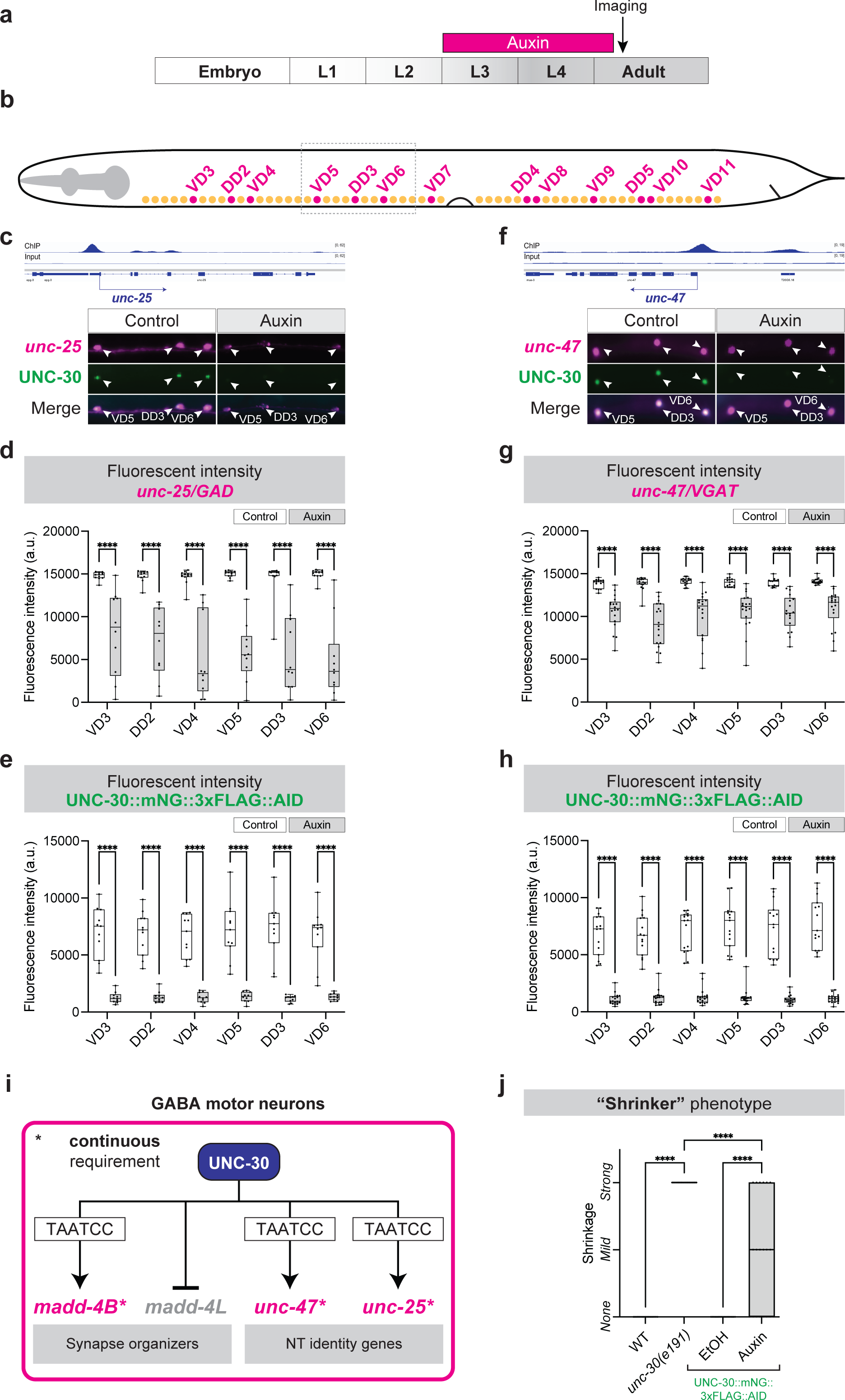
UNC-30 is required to maintain expression of GABA biosynthesis genes. (a) Auxin treatment timeline. Imaging occurred at day 1 adults. (b) GABAergic (magenta) and cholinergic (yellow) MNs. Dashed box depicts imaged area shown in (c,f). (c) Expression analysis of *unc-25(hpIs88 [unc-25p::mCherry)* and UNC-30 (*syb2344 [UNC-30::mNG::3xFLAG::AID])* in control (EtOH treated) and auxin-treated animals. Animals express TIR1 pan-somatically (*ieSi57 [Peft-3::TIR1::mRuby]*). White arrowheads: GABAergic MNs. UNC-30 ChIP-seq tracks on the *unc-25* and *unc-47* (f) loci. (d-e) Quantification of (d) *unc-25(hpIs88 [unc-25p::mCherry)* or (e) UNC-30(*syb2344 [UNC-30::mNG::3xFLAG::AID])* fluorescent intensity in GABAergic MNs. Two-way ANOVA followed by Sidak’s multiple comparison test. ****p<0.0001. Control: n=11, Auxin-treated=10. (f) Expression analysis of *unc-47(otIs565 [unc-47(fosmid)::SL2::H2B::mChopti)* and UNC-30(*syb2344 [UNC-30::mNG::3xFLAG::AID])* in control (EtOH treated) and auxin-treated animals. Animals express TIR1 pan-somatically (*ieSi57 [Peft-3::TIR1::mRuby]*). White arrowheads: GABAergic MNs. (g-h) Quantification of (g) *unc-47 (otIs565 [unc-47(fosmid)::SL2::H2B::mChopti))* or (h) UNC-30 (*syb2344 [UNC-30::mNG::3xFLAG::AID])* fluorescent intensity in GABAergic MNs. Two-way ANOVA followed by Sidak’s multiple comparison test. ****p<0.0001. Control: n=15, Auxin-treated=18. (i) UNC-30 is required to maintain *unc-25/GAD, unc-47/VGAT* and *madd-4S* expression in GABAergic MNs. (j) Quantification of the “shrinker” phenotype upon response. One-way ANOVA followed by Sidak’s multiple comparison test. ****p<0.0001. Wild-type: n=20, *unc-30(e191)*: n=20, Control: n=20, Auxin-treated: n=20. For panels d-e, g-h: Box and whisker plots show median, lower, and upper quartiles – whiskers represent minimum and maximum. Black circles depict values.

Whether UNC-30 is required at post-embryonic stages to maintain the expression of these critical determinants of GABAergic identity (e.g., *unc-25/GAD, unc-47/VGAT*) and function is not known. We again employed the AID system in late larval stages; this time assessing the effect of UNC-30 depletion on expression levels of *unc-25/GAD* and *unc-47/VGAT.* We observed a significant reduction in their expression levels in GABAergic MNs (**Figure 6a-h, Supplementary Figure 5**), suggesting that UNC-30 is not only required during early development to initiate expression of GABA biosynthesis genes, but also to maintain their expression during late larval stages (**Figure 6i**).

### UNC-30 is continuously required for normal touch response

Having established that UNC-30 is required at later stages to maintain expression of *unc-25/GAD*, *unc-47/VGAT* and *madd-4B/Punctin*, we then asked: is UNC-30 is also continuously required for normal animal behavior? Animals lacking unc-30 gene activity (homozygous null mutants) display a characteristic locomotory phenotype nicknamed “shrinker” ^45,55^, i.e., *unc-30* mutants hyper contract their body wall muscles in response to touch due to the lack of GABAergic MN inhibitory input to muscles. We indeed observed a striking and fully penetrant “shrinker” phenotype in *unc-30(e191)* mutants compared to control animals (**Figure 6j**). Importantly, auxin-mediated depletion of UNC-30 specifically at late larval/early adult stages also resulted in “shrinker” animals (**Figure 6j**). Because the auxin system does not fully eliminate UNC-30, as evidenced by quantification of UNC-30::mNG::3xFLAG::AID expression levels in individual GABAergic MNs (**Figure 5i**), the “shrinker” phenotype displays variable expressivity (none, mild, strong) upon auxin treatment (**Figure 6j**). We note that in the control (ethanol) condition, we observed no shrinkers, suggesting that tagging the endogenous *unc-30* gene with the mNG::3xFLAG::AID cassette does not result in detectable hypomorphic effects on locomotory behavior (**Figure 6j**). We therefore conclude that UNC-30 is continuously required for normal touch response.

### The dual role of UNC-30 in GABA MNs extends to other target genes

A handful of UNC-30 target genes are known to date, including *unc-25*/GAD, *unc-47*/VGAT*, pde-4/PDE4B, acy-1/ADCy9*, *oig-1, flp-11, flp-13,* and *ser-2* (**Table 1**)^15,16,47,51,56^. A unifying theme emerging from these studies is that UNC-30 acts as a transcriptional activator. However, our findings on *madd-4L* suggest a repressive role for UNC-30 in GABA MNs (**Figure 4**). We therefore sought to identify new UNC-30 target genes to determine whether the duality in UNC-30 function (activator and repressor) is broadly employed.

**Table 1.** Summary of genetically validated UNC-30/PITX targets in GABA MNs.

| Gene | Expression in GABA MNs* | UNC-30 binding (ChIP-Seq)# | *unc-30* dependency | Literature |
| --- | --- | --- | --- | --- |
| *unc-25/GAD* | Yes | Yes | Activated by UNC-30 | 16 |
| *unc-47/VGAT* | Yes | Yes | Activated by UNC-30 | 16 |
| *oig-1/one Ig* | Yes | Yes | Activated by UNC-30 | 15,46,47,51,85 |
| *flp-13/neuropeptide* | Yes | Yes | Activated by UNC-30 | 15,51,56 |
| *acr-14/AChR* | Yes | Yes | Activated by UNC-30 | 15 |
| *ser-2/tyramine receptor* | Yes | Yes | Activated by UNC-30 | 85 |
| *flp-11/neuropeptide* | Yes | Yes | Activated by UNC-30 | 85 |
| *pde-4/PDE4B* | Yes | Yes | Activated by UNC-30 | 51 |
| *acy-1/ADCY9* | Yes | Yes | Activated by UNC-30 | 51 |
| *madd-4S/Punctin* | Yes | Yes | Activated by UNC-30 | This study, <sup>50</sup> |
| *mab-9/Tbx20* | Yes | Yes | Activated by UNC-30 | This study |
| *aman-1/Man2b1* | Yes | Yes | Activated by UNC-30 | This study |
| *tsp-7/Cd63* | Yes | Yes | Activated by UNC-30 | This study |
| *nhr-49/Hnf4a* | Yes | Yes | Activated by UNC-30 | This study |
| *ilys-4* | Yes | Yes | No effect | This study |
| *nhr-40/NHR* | Yes | Yes | No effect | This study |
| *madd-4L/Punctin* | No | Yes | Repressed by UNC-30 | This study |
| *unc-53/NAV1* | No | Yes | Repressed by UNC-30 | This study |
| *glr-5/GRIK4* | No | Yes | Repressed by UNC-30 | This study |
| *nhr-19/NHR* | No | Yes | No effect | This study |
| *slo-2/KCNT* | No | Yes | No effect | This study |
| *gpd-2/GAPDH* | No | Yes | No effect | This study |
| *del-1/SCNN1* | No | Yes | No effect | This study |
| *cat-1/SLAC18A* | No | Yes | No effect | This study |
| *lin-11/LHX1* | No | Yes | No effect | This study |
| *srb-16* | No | No | No effect | This study |
| *unc-4/Uncx4.1* | No | No | No effect | This study |
| *ida-1/PTPRN2* | No | Yes | No effect | This study |

First, we identified putative *unc-30* targets by searching for UNC-30 binding peaks in genes that are normally expressed in GABA MNs^57,58^ (**Table 1**). In total, we tested six genes (*tsp-7/Cd63, aman-1/Man2b1, nhr-49/Hnf4a, mab-9/Tbx20, nhr-40/NHR, ilys-4*) by either generating new transgenic reporter animals (*nhr-49/Hnf4a, mab-9/Tbx20, nhr-40/NHR)*, or using available reporters (*tsp-7/Cd63, aman-1/Man2b1, ilys-4)*. Reporter expression for five of these genes (*tsp-7/Cd63, aman-1/Man2b1, nhr-49/Hnf4a, mab-9/Tbx20, nhr-40/NHR)* was significantly reduced in GABA MNs of *unc-30 (e191)* mutant animals (**Figure 7a-b**, **Table 1**). Because ChIP-Seq shows UNC-30 binding to all four of these genes (**Figure 7a-b**), it is likely that UNC-30 acts as a direct activator of *tsp-7/Cd63, aman-1/Man2b1, nhr-49/Hnf4a, nhr-40/NHR,* and *mab-9/Tbx20* transcription (**Figure 7d**).

**Figure 7:**
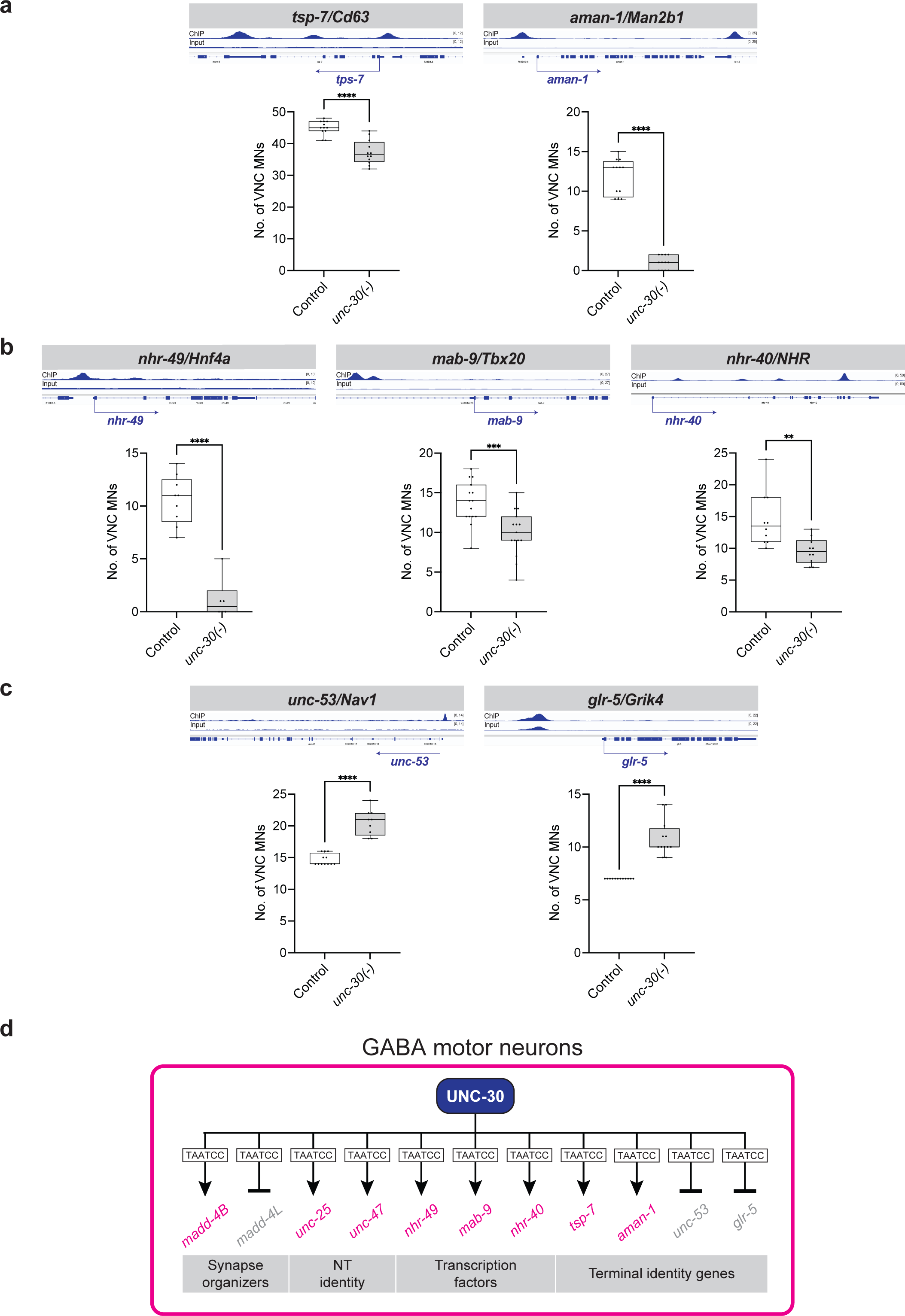
UNC-30 activates and represses different genes in GABAergic MNs. (a-c) Quantification of the number of GABAergic nerve cord MNs cord expressing *wdEx351 [tsp-7::GFP], sEx11477 [aman-1::GFP]), kasEx220 [nhr-49 (−803 to +58bp from ATG)::RFP, kasEx232 [mab-9 (−5569 to −3768bp from ATG)::RFP, kasEx214 [nhr-40(+938 to +1846bp from ATG)::RFP], hdIs1 [unc-53p::GFP],* or *icIs270 [glr-5::GFP]* in control and *unc-30(191)* animals. Stage: L4. Box and whisker plots show median, lower, and upper quartiles – whiskers represent minimum and maximum. Black circles depict values. Unpaired t-test with Welch’s correction. **p<0.002, ***p<0.0002, ****p<0.0001. Wild-type: n=10, *unc-30(e191)*: n=10. (d) Summary of UNC-30 targets in GABAergic motor neurons.

Next, we aimed to identify genes that, like *madd-4L*, are repressed by UNC-30. We searched for UNC-30 binding peaks in genes that are not expressed in GABA MNs, but instead are normally expressed in cholinergic nerve cord MNs (**Table 1**). In total, we tested 11 genes, for which transgenic reporter animals were available. Two (*unc-53*/NAV1 and *glr-5*/GRIK4) of the 11 reporters showed ectopic expression in GABA MNs of *unc-30 (e191)* mutant animals (**Figure 7c**, **Table 1**).

Altogether, our work identified nine new UNC-30 target genes; six are activated (*madd-4B/Punctin, tsp-7/Cd63, aman-1/Man2b1, nhr-49/Hnf4a, mab-9/Tbx20, nhr-40/NHR)* and three are repressed (*madd-4L/Punctin, unc-53*/NAV1, *glr-5*/GRIK4) by UNC-30 (**Figure 7d**). This analysis significantly expands the known repertoire of UNC-30/PITX target genes in the *C. elegans* nervous system (**Table 1 –** summary of UNC-30 targets), consolidating its previously known activator role and uncovering a putative repressive function.

## DISCUSSION

Here, we describe a molecular mechanism that coordinates two spatially separated processes critical for the function of chemical synapses. That is, NT biosynthesis in the presynaptic cell and NT receptor clustering at the postsynaptic cell. Using the *C. elegans* neuromuscular synapses as a model, we show that the terminal selector-type transcription factor UNC-30/PITX is required continuously to maintain expression of GABA identity genes (e.g., *unc-25/GAD, unc-47/VGAT*) in presynaptic GABAergic MNs, thereby ensuring GABA synthesis and release. In postsynaptic target muscle cells, UNC-30 acts non-cell autonomously to control clustering of GABA_A_Rs – the most prominent inhibitory NT receptors in animal nervous systems^38,39^. Mechanistically, we propose that UNC-30 directly regulates the production of MADD-4B, a secreted synapse organizer. Hence, UNC-30 coordinates GABA_A_R clustering on postsynaptic muscle cells with the acquisition and maintenance of GABAergic identity of presynaptic cells (**Figure 1c**, model 3), essentially safeguarding GABA neurotransmission. Further, this work advances our understanding of PITX transcription factor function in the nervous system by uncovering: a) a dual role (activator and repressor) for UNC-30 in GABA MNs, and b) multiple additional UNC-30 target genes. Altogether, our findings on UNC-30/PITX may help us understand the molecular mechanisms underlying the Axenfeld-Rieger syndrome and other genetic conditions caused by PITX gene mutations^40–42^.

### Transcriptional coordination of two spatially separated processes: NT biosynthesis in the presynaptic cell and postsynaptic NT receptor clustering

To enable synaptic output, the presynaptic neuron must synthesize, package, and release a specific NT, whereas the postsynaptic neuron must present cognate NT receptors. Whether and how these distinct and spatially separated processes are coordinated remains poorly understood. Here, we show that the conserved transcription factor UNC-30/PITX coordinates the process of GABA biosynthesis in *C. elegans* MNs with GABA_A_R clustering in postsynaptic target muscle cells. Through inducible protein depletion (**Figure 6**), we found that UNC-30 is required in late larval and adult stages to maintain expression of GABA biosynthesis genes (e.g., *unc-25/GAD, unc-47/VGAT*), consolidating its role as a terminal selector of GABA MN identity^16,17^. Further, UNC-30 acts directly to activate and maintain transcription of *madd-4B/Punctin,* a secreted synapse organizer necessary for GABA_A_ receptor clustering on target muscle cells^25^. Because UNC-30/PITX is continuously required in GABA MNs, this simple coregulatory strategy of *unc-25/GAD, unc-47/VGAT*, and *madd-4B/Punctin* by a terminal selector may ensure that key features of a functional synapse will continue to appear together throughout life (**Figure 1c**), ensuring continuous GABA neurotransmission. Hence, the presynaptic neuron will continue to synthesize and release GABA (ensured by continuous *unc-25/GAD* and *unc-47/VGAT* expression) and the postsynaptic neuron will constantly have the means to receive GABA via cognate receptor clustering (ensured by continuous m*add-4B/Punctin* expression). Because UNC-30 orthologs are expressed in planarian^59,60^, fly^61^, zebrafish^62^ and mouse nervous systems^63,64^, the coregulatory principle described here may be broadly applicable across species.

### Terminal selectors control synaptic connectivity

The only other known example of a terminal selector that operates in an analogous manner is UNC-3, the sole *C. elegans* ortholog of the COE (Collier/Olf/EBF) family of proteins^65^. In nerve cord cholinergic MNs, UNC-3 acts as a terminal selector, directly regulating scores of effector genes (e.g., ACh biosynthesis proteins, ion channels, neuropeptides)^66,67^. Like *unc-30*, *unc-3* is not expressed in *C. elegans* muscles. Yet, it is required for AChR clustering on muscle cells^37^. In cholinergic MNs, UNC-3 not only directly activates *madd-4B* (whose function in these cells is discussed in next section), but also *madd-4L*, which is required for AChR clustering ^37^. On the other hand, we find that UNC-30 activates *madd-4B* but represses *madd-4L* in GABA MNs, thereby ensuring expression of the appropriate *madd-4* isoform (*madd-4B*). Altogether, NT receptor clustering in *C. elegans* neuromuscular synapses is achieved by two different terminal selectors regulating, in distinct ways, the two isoforms of the same synapse-organizing molecule. UNC-3 activates both *madd-4B* and *madd-4L* in cholinergic MNs, whereas UNC-30 activates *madd-4B* but represses *madd-4L* in GABA MNs.

Besides NT receptor clustering, additional synaptic connectivity defects have been reported in MNs of *unc-3* and *unc-30* mutant animals^37,46,47,68,69^. Specifically, cholinergic MN input onto GABA MNs is disrupted in *unc-3* mutants^68^. In this case, UNC-3 controls *nrx-1/*neurexin, a synapse organizer necessary for AChR localization onto dendrites of GABA MNs^69^. Presynaptic specializations of cholinergic MNs onto muscle cells are also affected in *unc-3* mutants^37^, but the underlying mechanism remains unclear. On the other hand, UNC-30 has been implicated in *C. elegans* synaptic remodeling, as it is necessary to prevent premature synapse rewiring of DD neurons and aberrant synapse rewiring of VD neurons^46,47^. This is achieved by UNC-30 directly regulating OIG-1, a single Ig domain protein that functions as a synaptic organizer^46,47^. Consistent with a recent review^6^, this work and the aforementioned studies provide strong evidence for expanding the definition of terminal selector genes. That is, terminal selectors not only regulate effector genes critical for NT biosynthesis and neuronal signaling (e.g., ion channels, neuropeptides), but also control synaptic connectivity via the regulation of distinct synapse-organizing molecules.

It is tempting to speculate that mammalian terminal selectors may operate in an analogous manner. For example, the terminal selector of mouse spinal MNs, Isl1^70^, may control transcription of agrin, a MN-derived synapse organizer necessary for AChR clustering on mouse skeletal muscles^71^. Curiously, although the function of MADD-4/Punctin in *C. elegans* is reminiscent of mammalian agrin in mice, *agr-1 (*agrin ortholog) is not involved in *C. elegans* neuromuscular synapse formation^72^.

### Neuron type-specific regulation of synapse organizers

Synapse organizers are cell adhesion or secreted molecules that control synapse formation and/or maintenance^4,73^. Their adhesive and signaling properties mediate uni- or bidirectional signaling, enabling pre- and/or postsynaptic differentiation^74^. Understanding the spatiotemporal regulation of synapse organizers is important because synapses must be built at the right place and time. However, we know very little about the transcriptional mechanisms that control synapse organizer expression, in part because these molecules usually have multiple isoforms (e.g., neurexins, neuroligins, leukocyte common antigen-related receptor protein-tyrosine phosphatases [LAR-PTPRs], Agrin, MADD-4/Punctin)^75–77^. Multiple isoforms can be produced via either alternative RNA splicing or promoter usage. To date, substantial research has focused on the regulation of alternative splicing of synapse organizers (e.g., neurexin isoforms)^78^, leaving the transcriptional mechanisms underlying alternative promoter usage poorly understood. Our work uncovered a transcriptional mechanism for the spatial (neuron type-specific) regulation of two different isoforms (produced via alternative promoter usage) of the same synapse organizer (MADD-4/Punctin) in *C. elegans* neuromuscular synapses.

MADD-4L is only produced by cholinergic MNs^25,37^. Upon secretion, it promotes clustering of the levamisole-sensitive heteromeric ACh receptor (L-AChR) by an extracellular scaffold composed of LEV-10 (LEVamisole resistant-10), LEV-9 and OIG-4 (One ImmunoGlobulin domain-4) ^25,30–32^. On the other hand, MADD-4B is produced by both cholinergic and GABAergic MNs. At GABAergic neuromuscular synapses, MADD-4B promotes GABA_A_R clustering on muscle cells through binding to NLG-1/ neuroligin and activation of UNC-40/DCC signaling^26–28^. At cholinergic neuromuscular synapses, MADD-4B inhibits the attraction of GABA_A_ receptors by MADD-4L^25^. Hence, spatial (neuron type-specific) regulation of MADD-4 isoform expression is crucial for the formation and function of excitatory (ACh) and inhibitory (GABA) synapses in *C. elegans.* Our previous work identified UNC-3 as a critical activator of both *madd-4* isoforms in cholinergic MNs^37^. Here, we show in GABA MNs that UNC-30 controls the two *madd-4* isoforms in opposite ways; it provides direct and positive input to the *madd-4B* promoter and negative input to the *madd-4L* promoter, thereby ensuring proper GABA_A_ receptor clustering on target muscle cells.

### Advancing our understanding of PITX gene function in the nervous system

In humans, *PITX* gene mutations cause various congenital defects and cancer^42^. *Pitx* genes belong to the PAIRED (PRD) class of highly conserved homeobox genes; vertebrates have three *Pitx* genes and invertebrates have one. In mice, *Pitx* genes play critical roles in the development of the nervous system, craniofacial structures, and limbs [reviewed in ^42^]. *Pitx2* and *Pitx3* are expressed in discrete cell populations of the mouse midbrain and spinal cord, and their expression persists into adult stages^63,64,79^. *Pitx3* is essential for the survival of dopaminergic neurons of the substantia nigra, a key cellular substrate of Parkinson’s disease^64^. Importantly, human mutations in *PITX2* or *PITX3* affect eye development^42,80^. *PITX2* mutations cause Axenfeld-Rieger syndrome, a primarily eye disorder that also affects the craniofacial and cardiovascular systems, whereas PITX3 mutations are associated with congenital cataracts^40,41^.

Mechanistically, functional assays showed that human *PITX2* and *PITX3* gene mutations result in reduced transcriptional activity^81–83^. However, the transcriptional targets of PITX proteins remain poorly defined and whether they act as transcriptional activators and/or repressors is not well defined. Our study contributes to these knowledge gaps in three ways. First, we identify nine new UNC-30 target genes (*madd-4B/PunctinB, madd-4L/PunctinL, mab-9/Tbx20, nhr-49/Hnf4a, nhr-40/NHR, tsp-7/Cd63, aman-1/Man2b1, unc-53*/NAV1, *glr-5*/GRIK4), significantly expanding the list of PITX targets in the nervous system (**Table 1**). Second, consistent with its previously described direct mode of activation of genes involved in GABA biosynthesis and neuronal rewiring ^16,46,47,51^, our mutational analysis and available ChIP-Seq data indicate that UNC-30 primarily acts directly to control expression of these eight genes. Last, we found that six genes are activated (*madd-4B, tsp-7, aman-1, mab-9, nhr-40, nhr-49*) and three are repressed (*madd-4L, unc-53, glr-5*) by UNC-30. Because it binds directly to the *cis*-regulatory region of most of these genes, we propose that, in GABA MNs, UNC-30 acts as a direct activator and direct repressor of distinct sets of genes. Intriguingly, a similar dual role for UNC-30 has recently been described in *C. elegans* glia^84^.

### Limitations of this work

Future studies are needed to dissect the molecular mechanism underlying the dual role of UNC-30 in GABA MNs. It is likely that cooperation with distinct transcription factors shifts its transcriptional activity from an activator to a repressor. Candidates include LIN-39/HOX, a known transcriptional activator in GABA MNs^85^, and UNC-55/NR2F, a known transcriptional repressor in these cells^51,56^. Another limitation relates to maintenance of GABA_A_R clustering. Although we showed that UNC-30 is required to maintain *madd-4B* transcription in late larval/early adult stages, it remains unknown whether inducible UNC-30 depletion at these stages affects maintenance of GABA_A_R clustering. Last, our work is focused on neuromuscular synapses. Notably, Punctin (ADAMTSL3) and other secreted synapse organizers (e.g., cerebellins, pentraxins, Sema3F, BDNF) are expressed in the mammalian brain ^36,86,87^ ^35^. Hence, similar coregulatory strategies to the one described here may operate in neuron-neuron or neuron-glia synapses in the central nervous system.

## MATERIALS AND METHODS

### C. elegans strains

Worms were grown at 15°C, 20°C or 25°C on nematode growth media (NGM) plates seeded with bacteria (*E.coli* OP50) as food source. All *C. elegans* strains used in this study are listed in **Supplementary file 1.**

### Generation of transgenic reporter animals

Reporter gene fusions for *cis*-regulatory analysis were made using either PCR fusion or Gibson Assembly Cloning Kit (NEB #5510S) ^88^. Targeted DNA fragments were fused (ligated) to *tagrfp* or *gfp* coding sequence, followed by *unc-54 3’ UTR.* Mutations of UNC-30 binding sites were introduced via PCR mutagenesis. The product DNA fragments were either injected into young adult *pha-1(e2123)* hermaphrodites at 50ng/µl using *pha-1* (pBX plasmid) as co-injection marker (50 ng/µl) and further selected for survival, or injected into young adult N2 hermaphrodites at 50ng/µl (plus 50ng/µl pBX plasmid) using *myo-2::gfp* as co-injection marker (3 ng/µl) and further selected for GFP signal. Primer sequences used for reporter construct generation are provided in **Supplementary file 2**.

### Targeted genome engineering

CRISPR/Cas9 genome editing was performed by SunyBiotech following standard procedures^89^. The *unc-30* endogenous reporter allele *syb2344 [unc-30::mNG::3xFlag::AID]* was generated by SunyBiotech via CRISPR/Cas9 by inserting the *mNG::3xFLAG::AID* cassette immediately before the *unc-30* termination codon. The endogenous *madd-4L* reporter allele syb624 *[2xNLS::mScarlet::SL2::madd-4L]* was generated by inserting the *2xNLS::mScarlet::SL2* cassette immediately after the ATG of *madd-4L*. The endogenous *madd-4B* reporter allele *syb623 [2xNLS::mScarlet::SL2::madd-4B]* was generated by inserting the *2xNLS::mScarlet::SL2* cassette immediately after the ATG of *madd-4B*. The endogenous *madd-4B* reporter allele *syb3561 [2xNLS::mScarlet::SL2::madd-4B* ^Δ506^ ^bp^] was generated by creating a 506 bp-long deletion (−1433bp to −927bp from the *madd-4B* ATG) in the background strain carrying the endogenous *madd-4B* reporter allele *syb623 [2xNLS::mScarlet::SL2::madd-4B]*.

### Temporally controlled protein degradation

AID-tagged proteins are conditionally degraded when exposed to auxin in the presence of TIR1^53,54^. Animals carrying auxin-inducible alleles of *unc-30 (syb2344[unc-30::mNG::3xFLAG::AID]) IV* were crossed with *ieSi57* animals that express TIR1 pansomatically. Auxin (indole-3-acetic acid [IAA], Catalog number A10556, Alfa Aesar) was dissolved in ethanol (EtOH) to prepare 400 mM stock solutions which were stored at 4°C for up to one month. NGM agar plates were poured with auxin or ethanol added to a final concentration of 4 mM and allowed to dry overnight at room temperature. Plates were seeded with OP50 bacteria. To induce protein degradation, worms of the experimental strains were transferred onto auxin-coated plates and kept at 20°C. As a control, worms were transferred onto EtOH-coated plates instead. Auxin solutions, auxin-coated plates, and experimental plates were shielded from light.

### Microscopy

For Figure 2, young adult *C. elegans* were mounted on 2% agarose (w/v in water) dry pads immersed in 2% polystyrene bead (0.1 mm diameter, Polyscience, 00876-15) diluted in M9 buffer. Images were taken using a Nikon-IX86 microscope (Olympus) equipped with an Andor spinning disk system (Oxford Instruments), a 60x/NA1.42 oil immersion objective and an Evolve EMCCD camera. For each animal (Figure 2), an image of the dorsal nerve cord at the first quarter of the worm was acquired as a stack of optical sections (0.2 μm apart). The Pearson’s coefficient was calculated as described^26^. For the remaining figures, worms were anesthetized using 100mM of sodium azide (NaN_3_) and mounted on a 4% agarose pad on glass slides. Images were taken using an automated fluorescence microscope (Zeiss, Axio Imager.Z2). Several z-stack images (each ∼1 µm thick) were acquired with a Zeiss Axiocam 503 mono using the ZEN software (Version 2.3.69.1000, Blue edition). Representative images are shown following max-projection of 1-8 µm Z-stacks using the maximum intensity projection type. Image reconstruction was performed using Image J/FIJI software^90^.

### MN identification

MNs were identified based on a combination of the following factors: i) co-localization with fluorescent markers with known expression pattern, ii) invariant cell body position along the ventral nerve cord, or relative to other MN subtypes, iii) MN birth order, and (iv) number of MNs that belong to each subtype.

### Fluorescence Intensity (FI) Quantification

To quantify FI of individual MNs in the VNC, images of worms from different genetic backgrounds were taken with identical parameters through full-thickness z-stacks that cover the entire cell body. Image stacks were then processed and quantified for FI via FIJI. The focal plane in Z-stacks that has the brightest FI was selected for quantification. Background signal was minimized by using FIJI’s background subtraction feature (rolling ball at 50 pixels). Cell outline was manually selected, and FIJI was used to quantify the FI and area to get the mean value for FI.

### Statistical analysis and Reproducibility

For quantification, box and whisker plots were adopted to represent the quartiles in graph. The box includes data points from the first to the third quartile value with the horizontal line in box representing the median value. Upper and lower limits indicate the max and min, respectively. Unpaired t-test with Welch’s correction was performed and p-values were annotated. Visualization of data and p-value calculation were performed via GraphPad Prism Version 9.2.0 (283). Each experiment was repeated twice.

### Immunofluorescence staining

For Figure 2, immunofluorescence staining was performed as described^26^. Images were acquired using a Leica 5000B microscope equipped with a spinning disk CSU10 (Yokogawa) and a Coolsnap HQ2 camera.

## ACKNOWLEDGEMENTS

We thank the *Caenorhabditis* Genetics Center (CGC), funded by NIH Office of Research Infrastructure Programs (P40 OD010440), for providing strains. We thank members of the Kratsios lab (Mira Antonopoulos, Jayson Smith, Filipe Marques, Anthony Osuma, Manasa Prahlad) for providing feedback, and Yihan Chen and Jihad Aburas for technical assistance. This work was supported as follows: E.C.: F31NS124277, T32GM007183, 5R25GM109439; M.M.: fellowship from the French Ministry of Research; JLB: ERC_Adg C.NAPSE #695295, LABEX CORTEX (ANR-11-LABX-0042) of University Lyon 1, within the program “Investissements d’Avenir” (ANR-11-IDEX-0007); B.P.L.: INSERM support; P.K.: R01 NS118078 (NIH) and R01 NS116365 (NIH).

## AUTHOR CONTRIBUTIONS

Conceptualization: E.C.,M.M.,B.P.L,J.L.B.,P.K.; Methodology: E.C.,M.M.,M.C.; Investigation: E.C.,M.M.,M.C.; Formal analysis: E.C., M.M.,M.C; Visualization: E.C., M.M.,B.P.L,P.K.; Funding acquisition: J.L.B.,P.K.; Writing original draft. E.C.,B.P.L.,P.K.; Writing - review and editing: E.C.,M.M.,M.C.,B.P.L,J.L.B.,P.K.; Supervision: B.P.L,J.L.B.,P.K.

---

**Supplementary Figure 1:**
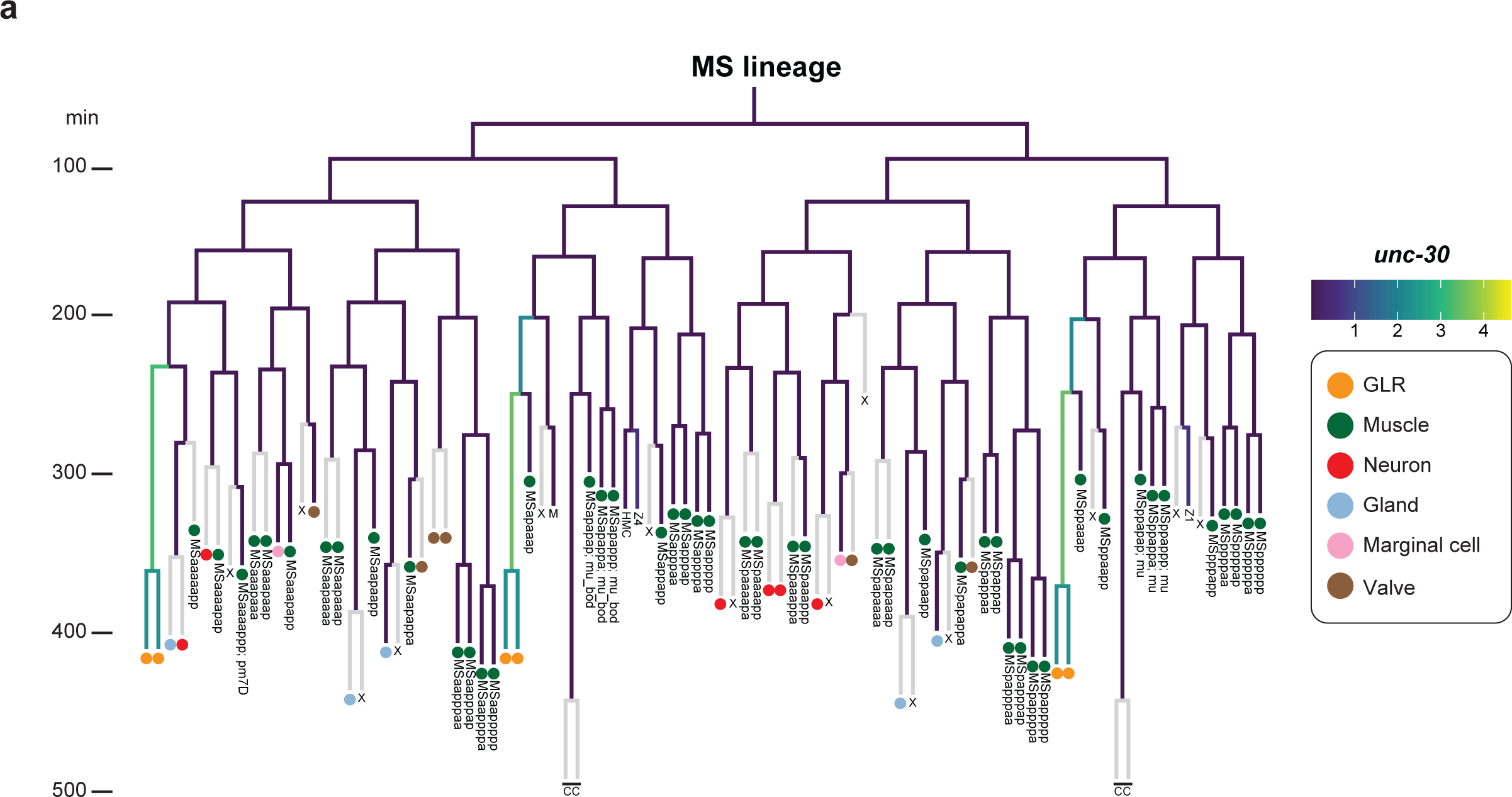
*unc-30* is not expressed in *C. elegans* body wall muscle cells. (a) Lineage of the blast cell MS. Gradient depicts *unc-30* mRNA expression, while grey color depicts cells that were not characterized for *unc-30*. Single-cell RNA-Sequencing data were analyzed from ^92^.

**Supplementary Figure 2:**
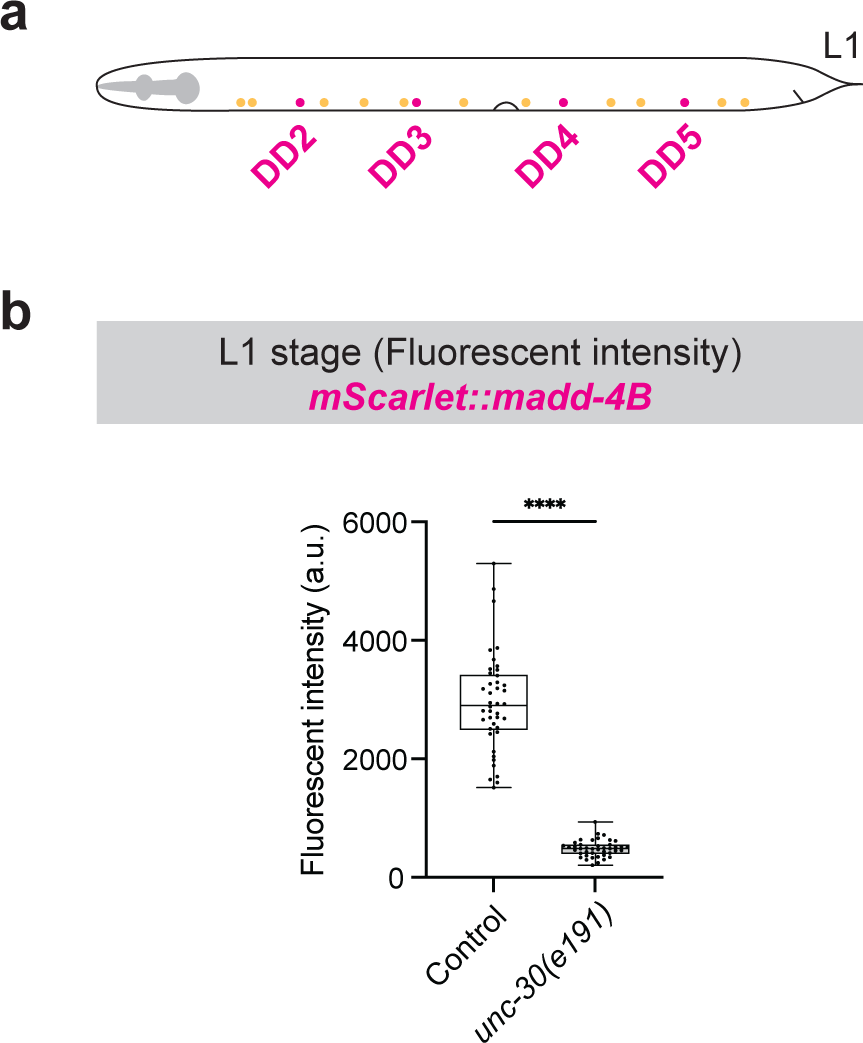
UNC-30 directly activates *madd-4B* in GABAergic MNs at larval stage 1. (a) Schematic of GABAergic (magenta) and cholinergic (yellow) nerve cord MNs at L1. MNs depicted are embryonically born. (b) Quantification of *madd-4B(syb623 [2xNLS::mScarlet::SL2::madd-4B])* fluorescent intensity in GABAergic MNs at L1 in control and *unc-30(191)* animals. Animals also carry a homozygous cholinergic Mn reporter *(otIs354[cho-1(fosmid)::SL2::YFP::H2B]).* Box and whisker plots show median, lower, and upper quartiles – whiskers represent minimum and maximum. Black dots depict values. Unpaired t-test with Welch’s correction. ****p<0.0001. Control: n=45 MNs, *unc-30(e191)*: n=45 MNs.

**Supplementary Figure 3:**
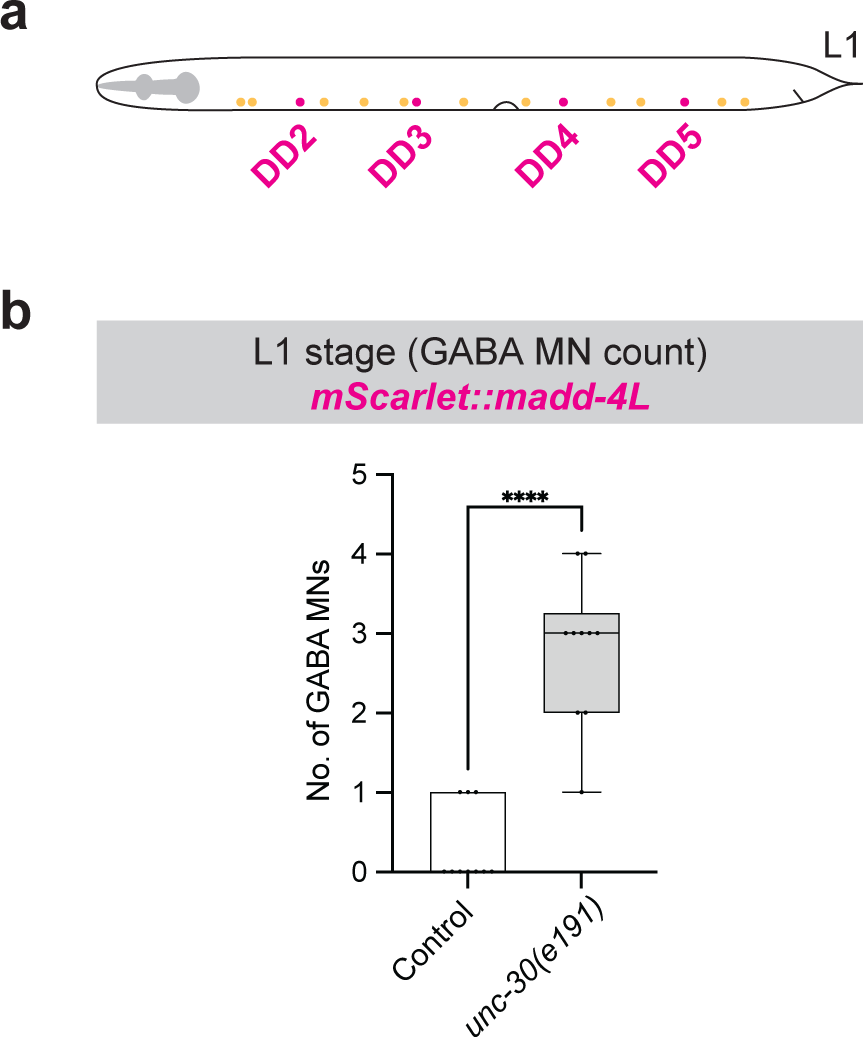
UNC-30 represses *madd-4L* in GABAergic DD motor neurons at larval stage 1. (a) Schematic of GABAergic (magenta) and cholinergic (yellow) MNs at L1. MNs depicted are embryonically born. (b) Quantification of GABAergic MNs expressing *madd-4L(syb624 [2xNLS::mScarlet::SL2::madd-4L])* at larval stage1 in control and *unc-30(191)* animals. Animals carry a cholinergic MN reporter *(otIs354 [cho-1(fosmid)::SL2::YFP::H2B]).* Box and whisker plots show median, lower, and upper quartiles – whiskers represent minimum and maximum. Black circles depict values. Unpaired t-test with Welch’s correction. ****p<0.0001. Wild-type: n=45, *unc-30(e191)*: n=45.

**Supplementary Figure 4:**
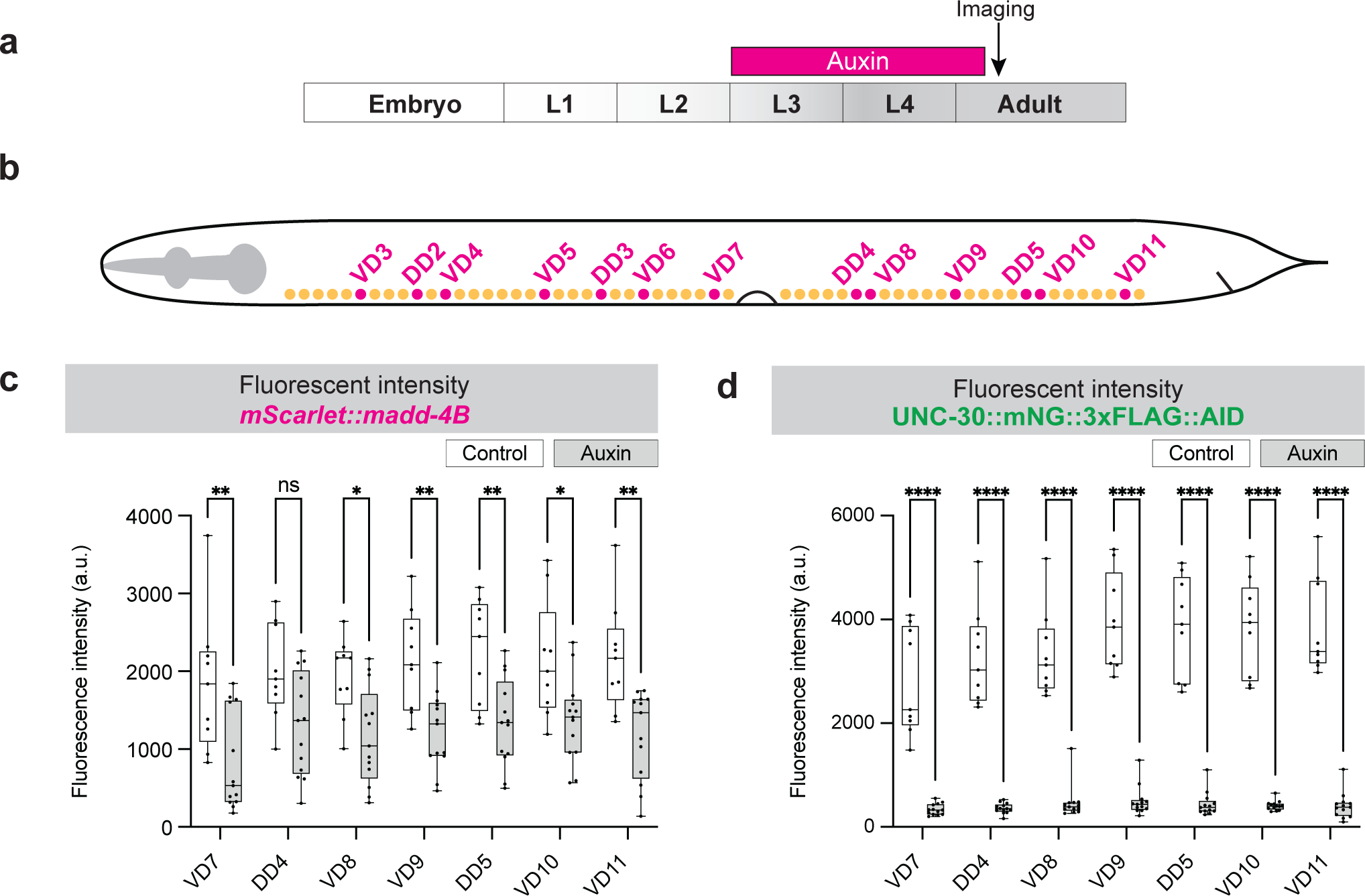
UNC-30 is required to maintain *madd-4B* expression in GABAergic MNs. Related to Figure 5. (a) Schematic of auxin treatment timeline. (b) Schematic of GABAergic (magenta) and cholinergic (yellow) nerve cord MNs. (c-d) Quantification of *madd-4B (syb623[2xNLS::mScarlet::SL2::madd-4B])* or UNC-30(*syb2344 [UNC-30::mNG::3xFLAG::AID])* fluorescent intensity in individual posteriorGABAergic MNs. Animals express TIR1 pan-somatically (*ieSi57 [Peft- 3::TIR1::mRuby]*). Box and whisker plots show median, lower, and upper quartiles – whiskers represent minimum and maximum. Black circles depict values. Two-way ANOVA followed by Sidak’s multiple comparison test. ****p<0.0001. Control: n=11, Auxin-treated=10.

**Supplementary Figure 5:**
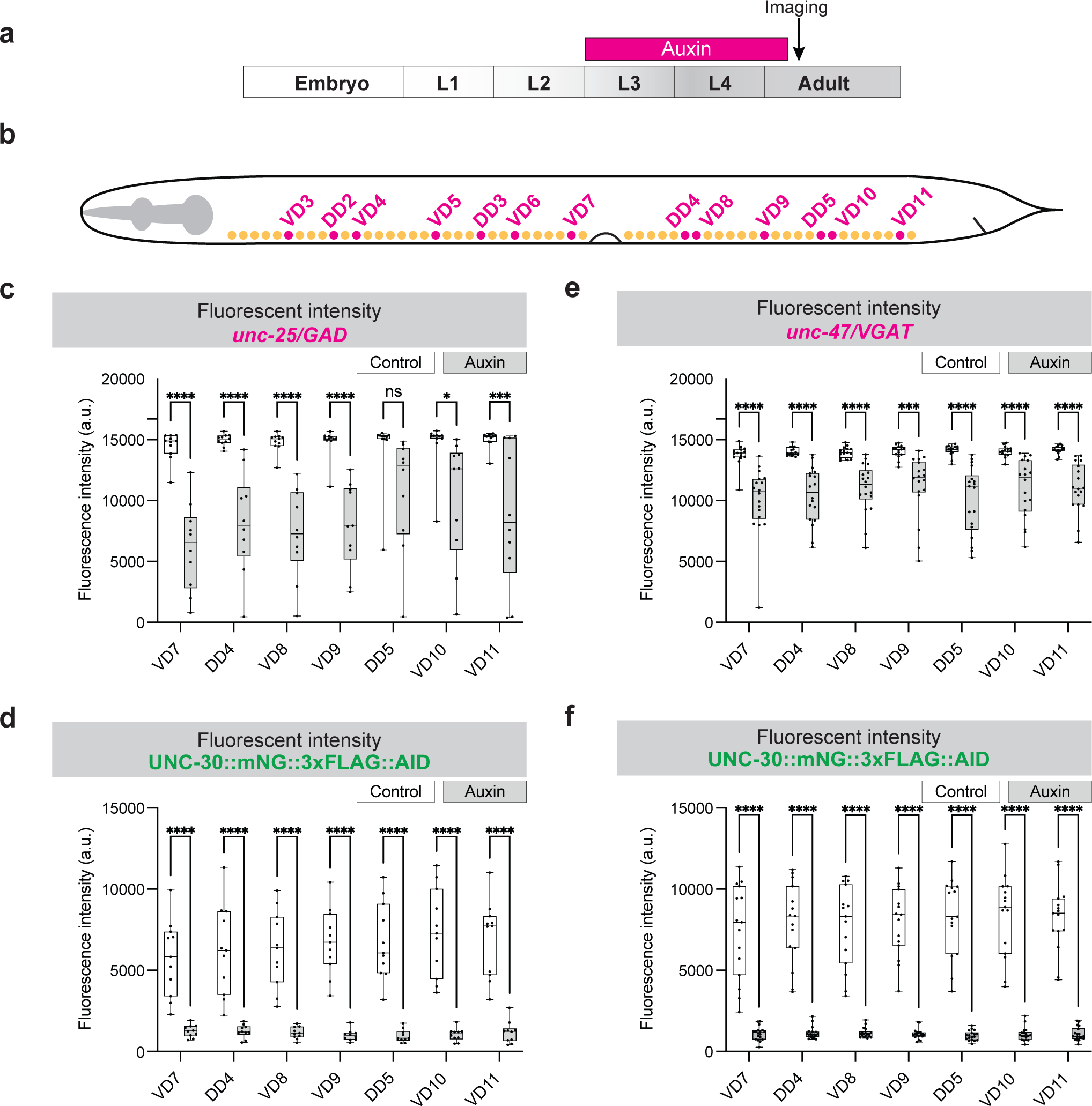
UNC-30 is required to maintain expression of GABA biosynthesis genes in MNs. Related to Figure 6. (a) Schematic of auxin treatment timeline. (b) Schematic of GABAergic (magenta) and cholinergic (yellow) MNs. (c-d) Quantification of *unc-25 (hpIs88 [unc-25p::mCherry)* or UNC-30 (*syb2344 [UNC- 30::mNG::3xFLAG::AID])* fluorescent intensity in individual posterior GABAergic MNs. Animals express TIR1 pan-somatically (*ieSi57 [Peft-3::TIR1::mRuby]*). Box and whisker plots show median, lower, and upper quartiles – whiskers represent minimum and maximum. Black circles depict values. Two-way ANOVA followed by Sidak’s multiple comparison test. ****p<0.0001. Control: n=11, Auxin-treated=10. (e-f) Quantification of *unc-47 (otIs565 [unc-47(fosmid)::SL2::H2B::mChopti))* or UNC-30 (*syb2344 [UNC-30::mNG::3xFLAG::AID])* fluorescent intensity in posterior GABAergic MNs. Animals express TIR1 pan-somatically (*ieSi57 [Peft-3::TIR1::mRuby]*). Box and whisker plots show median, lower, and upper quartiles – whiskers represent minimum and maximum. Black circles depict values. Two-way ANOVA followed by Sidak’s multiple comparison test. ****p<0.0001. Control: n=15, Auxin-treated=18.

**Supplementary File 1:** List of *C. elegans* strains used in this study.

**Supplementary File 2:** List of primers used in this study.

## Notes for Table 1

Asterisk (*) indicates expression based on single-cell RNA-Seq data from^57,58^

\# indicates UNC-30 binding based on ChIP-Seq data from^51^

